# Poka: a necro-robot beetle with a measured payload ratio of 6847 %

**DOI:** 10.1101/2024.11.27.625760

**Authors:** Yordan Tsvetkov, Parvez Alam

**Affiliations:** School of Engineering, The University of Edinburgh, Robert Stevenson Road, Edinburgh EH9 3FB, UK

**Keywords:** Necro-robot, Necrobot, Rhinoceros beetle, Mechanical design, Robotics, Bionic engineering

## Abstract

This paper is concerned with the design, manufacture and validation of ‘Poka’, a novel millimetre-scale necro-robot aimed at bridging the performance gap between miniature robots and insects. To create Poka, we use the exoskeleton of a deceased five-horned rhinoceros beetle (*Eupatorus gracilicornis*) as a mechanical chassis, which is mechatronically functionalised to enable ambulation. When comparing the payload ratio, *PR*, of Poka against reported values of the rhinoceros beetle *Xyloryctes thestalus*, we find that Poka’s *PR* is more than 2-fold higher, reaching a measured maximum of 6847% (i.e. 68.47 times its own body weight). The specific power at maximum payload, *P*_*s,t*_, is nevertheless of the same order of magnitude in both *Xyloryctes thestalus* (0.21 W/kg) and Poka (0.28 W/kg). Poka’s highest average speed, 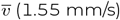 is achieved at a *PR* = 2739%, after which it progressively decreases with increasing payload ratio, reaching its minimum 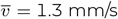 at maximum payload ratio. When comparing Poka’s maximum measured *PR* of 6847% against those of sixteen other ambulating robots, we find that Poka’s *PR* far exceeds that of any other robot to date, the highest being otherwise from SuperBot who has a *PR* = 530%. Poka’s payload ratio is therefore the highest robot payload ratio recorded to date and we attribute this to (a) the use of the beetle body as a natural composite chassis with high specific properties, and (b) the additive manufacture of bionic beetle parts using low density but stiff polylactic acid, designed with structurally stable geometries.

## INTRODUCTION

Small-scale robotics is a field of growing interest due to the potential benefit in a plethora of diverse applications including in machinery and civil infrastructure inspection, in complex medical operations, in reconnaissance, and in high-precision manufacturing ^1,2^. The design of miniature mechanical systems nevertheless presents significant engineering challenges, particularly in terms of locomotion, and in the incorporation of small-scale electronics, required for autonomous operation ^3^. Biomimetic robots, drawing design inspiration from biological concepts and mechanisms, show promise in addressing these challenges. Insects for example are potential sources of inspiration in robotics, due to their high functionality at small length scales ^4–6^, having already incentivised the development of small-scale technologies such as adhesives ^7^, surface coatings ^8^, and sensors ^9^. Beetles and cockroaches show promise in impacting the mobility design of small-scale robots ^6,10^. This is because they have the capacity to maintain structure and posture when bearing high payloads ^11,12^, even when on steep gradients ^13^. Insects have numerous mechanical advantages at the milimetre length scale. Insect exoskeletons (sclerites) are of high stiffness and strength ^14^, while insect legs combine hardened chitin sections with soft intersegmental membranes ^15^, to produce segmentally stiff composite limbs with controllable compliance, which can be leveraged for passive perturbance rejection.

Insects exhibit a variety of locomotive forms including walking ^16–18^, jumping ^19–21^, swimming ^22,23^, skating ^24^, flying ^25–27^, crawling ^28^, and climbing ^29,30^, in most cases their primary locomotor structures being legs and wings. Insect locomotion has been researched from a broad range of perspectives including kinematics ^31,32^, materials ^33–36^ and mechanics ^37^. Beetle locomotion has been of particular interest as there are several examples of beetle species with large bodies that can be supported by their legs or lifted by their wings. Kram for example, evaluated the energy costs per weight of locomotion under load of the rhinoceros beetle species *Xyloryctes thestalus*, which was recorded to have a payload ratio of up to 3000%, carrying therefore, up to 30 times it’s own weight ^38^. Importantly, Kram further reported in this paper, that as the payload weight increased, the rate of energy consumption per total mass decreased, suggesting that beetle locomotion becomes significantly more efficient as a function of increased loading. This is presumably due to the structural design and composition of beetle legs ^39^, the joints of which are interlocked ^40^ multi-level structures that enhance mechanical performance in high-load bearing beetles (such as dung beetles ^41^). In addition, tribological properties such as friction are enhanced by the combination of claw grip ^42^ and fibrillar foot pads ^43,44^.

Insects possess locomotive capabilities which could be of interest to robotics researchers. They are able to carry multiple times their own weight, while their energy efficiency increases with heavier loads. Ramdya et al. ^45^ used computational methods to characterise the gait of *Drosophila melanogaster* by mapping leg trajectories from real-world experiments, and encoding them into a simplified model that considers the leg contact phases. Although a bipod gait where only two legs touch the ground at a time is the fastest possible gait on a level flat surface, Ramadya and co-workers found that tripod gaits were faster on banked surfaces, highlighting the importance of stabilisation for speed retention on gradients. In a tripod gait, two sets of three legs move synchronously with one another, which indicates that it is possible to achieve similar movements using only one or two rotary motions. While insect legs possess a multitude of degrees of freedom, they usually use them in coupled bipod or tripod gaits. This suggests that the underlying motions can be described with fewer degrees of freedom, as the trajectories of their legs can be fully described within the sagittal frame, implying that only two degrees of motion would be needed to enable similar motion.

The combination of energy efficiency coupled with simplified motivity has generated a wave of interest in insect inspired robotics. Birkmeyer et al. ^3^ for example describe the design of DASH, a small hexapedal robot with running velocities up to 15 body lengths per second (0.15 m/s). The design of DASH is biomimetic, employing a sprawled posture and an alternating tripod gait commonly seen in insects ^45^, as well as a compliant body to absorb impact energy on falling. In DASH, the challenge of manufacturing the linkages for the transformation of the rotary motion of the motor to the legs was handled by flexures. These were manufactured by cutting channels into a polymer composite to create single-part components comprising many flexural joints. Jayaram et al. ^46^ designed HAMR-JR, a quadrupedal robot weighing only 0.32 g, thus smaller than DASH. HAMR-JR ambulates using four legs rather of six, but is manufactured using SCM-Fabrication processes, similarly to DASH. Unlike DASH nevertheless, HAMR-JR is a tethered robot. Instead of using conventional DC motors, HAMR-JR employs piezoelectric actuators, which are low-weight and capable of high-frequency movement. The dynamic characteristics of the flexures and the frequency response of the actuators were characterised to derive the control strategies for HAMR-JR, considering the running speeds for trotting and pronking (stotting). Di Canio et al. ^47^ used the flexible end (tarsus) of dung beetle legs to inspire the design of legs with enhancements for ambulation over rough substrates. The tarsus consists of small rigid segments connected by compliant joints with a low range of motion. A flexible tarsus reduces the impact forces experienced on contact with the ground, and furthermore reduces the energy consumed for movement as a function of weight. Robotic design conjoining both the morphological and kinematic characteristics of dung beetles has proven to enable improved performance over conventional hexapod robots ^48^, specifically to enable increased step length without over-burdening the motors, longer reach with reduced torque requirements, and reduced acceleration requirements during walking. In conclusion, we can state that there is precedent for using the anatomy of insects, namely cockroaches and beetles, as inspiration for high-performing biomimetic designs. This has been demonstrated for both centimetre- and millimetre-scale robots, the smallest of which can carry its own power supply is DASH, which is approximately twice as long and 7 times as heavy as a five-horned rhinoceros beetle.

The repurposing of animal body parts of interest, since animal body parts are readily available pre-formed structures that are geometrically suitable for robot construction. They are renewable, sustainable and biodegradable, and have been “field tested” by the animal. There are nevertheless still only a few examples of bio-engineering hybrids such as cyborg insects ^49–54^ and necro-robots (or necrobots) ^55,56^. Yap et al. ^55^ demonstrated a gripper manipulator that makes use of the exoskeleton of a deceased wolf spider. The gripper takes advantage of the pre-formed nature of the biological component for simplified manufacturing, requiring only the insertion of a sealed hypodermic needle in the spider’s prosoma. The resulting gripper was characterised by its ability to manipulate irregular objects 1.34 times heavier than itself, its biodegradability, and its unobtrusiveness in a natural environment. The complexities in using a biological robotic component were evaluated and described, with the primary limitation of the gripper being its short life cycle. Due to the evaporation of moisture from the joints of the legs, the gripper can become brittle and is thus susceptible to mechanical fracture, ultimately leading to a functional duration of only two days. The problem of biological degradation in the development of necro-robots has been discussed by Jørgensen and co-workers ^56^, who considered specifically, appropriate mechanisms by which means dog skulls could be rearticulated at the manidble symphysis for use as necro-robotic end effectors. While there is an engineering argument to be made for necro-robotics (or, necrobotics), it can also be used advantageously in other fields such as paleobiology, an example of which includes the reverse engineering of a necro-robot from a stem amonite (*Orobates pabsti*), which helped provide insights into its locomotive abilities ^57^.

To bridge the performance gap between miniature robots and insects, in this paper we propose using the body of a deceased five-horned rhinoceros beetle (*Eupatorus gracilicornis*) as a pre-formed chassis, mechatronically modified for locomotion. Using the body of the beetle will allow us to leverage the chitin exoskeleton, to enable both the rapid and cheap production of milimetre-scale robots, without the need for specialised manufacturing equipment, whilst taking advantage of a lightweight yet strong exoskeletal structure. Our aim is to create a necro-robot beetle, hereafter referred to as ‘Poka’, with an extreme payload-to-weight ratio of *at least* 3000% (i.e. 30 times its own weight), which will be comparable with the capabilities of living rhinoceros beetles ^38^.

## RESULTS AND DISCUSSION

### Morphometric characterisation

Figure 1 shows representative 3D scans of *Eupatorus gracilicornis* from lateral, dorsal, anterial and isometric perspectives. This is a simplified model representing a volume offset of 0.5 mm from the scanned mesh to account for scanning noise. Our key morphometric measurements for Poka, were extracted from this model. Poka was generally of a similar size and profile to male beetles within a measured batch (n = 13), with mean values including the standard deviations about the mean (±SD) and the coefficients of variation (CoV), as follows: mean total length of beetles was 71.6 mm (SD ± 3.4 mm, CoV 4.8 %), and the mean beetle length without the horn was 55 mm (SD ± 2.1 mm, CoV 3.8 %). The beetle selected to demonstrate the conversion to a necro-robot had the dimensions show in Table 1. The total length for this beetle was within one standard deviation of the mean total length for the sample set. The internal components of this necro-robot demonstrator would be expected to fit into the volume spaces shown in this table.

**Table 1:** Dimensions of the beetle body intended for use as the necro-robot demonstrator, Poka.

|  |  |
| --- | --- |
| Length, without horn [mm] | 58.15 |
| Length, with horn [mm] | 74.45 |
| Abdomen internal volume, length [mm] | 31.40 |
| Abdomen internal volume, width [mm] | 38.46 |
| Abdomen internal volume, height [mm] | 21.33 |
| Pronotum internal volume, diameter [mm] | 16.00 |
| Pronotum internal volume, length [mm] | 11.90 |

**Figure 1:**
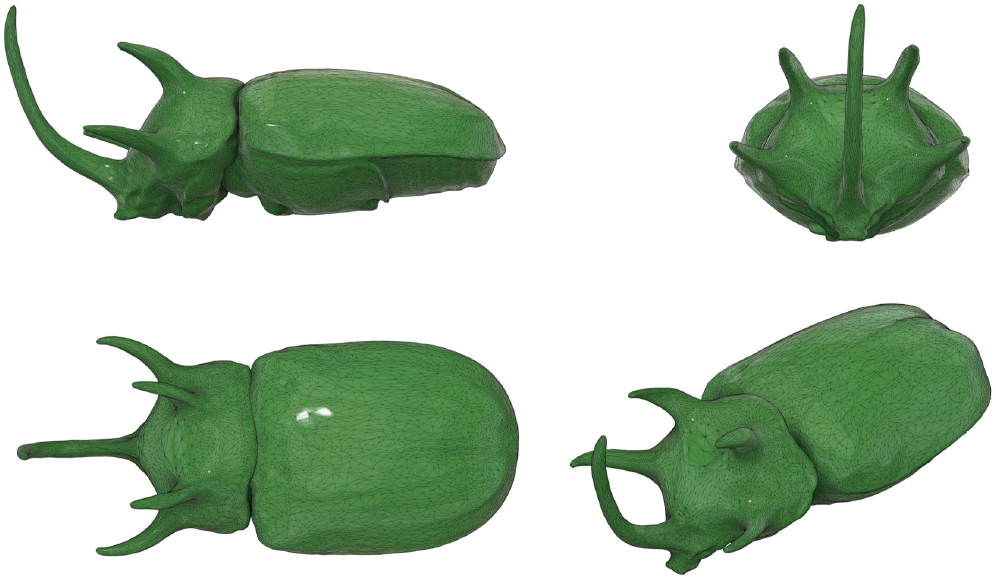
Visualisation of obtained 3D scan in Fusion 360 (Autodesk, San Francisco, CA.

### Necro-robot design

Aside from the dimensional constraints imposed by the natural internal geometry of a demonstrator beetle, the design of Poka posed the additional coupled load-bearing/locomotion challenge. Poka would need to be able to bear high relative payloads and as such, we aim to minimise friction of the internal moving components to maximise the transmission of mechanical energy, thus optimising the transmission of electrical to mechanical energy. This in turn has the beneficial effect of reducing both power generation and application losses. To address the coupled load-bearing/locomotion challenge, we considered the following: (1) component configuration where the total number of cams, cam-linkages, motors, and supporting elements (and their sizes) required mapping prior to selection (2) cam design based on both dimensional constraints and low-friction functionality (3) linkage design based on the both cam design and calculations for motion generation (4) selection of appropriate mechanisms for actuation and (5) the design of a ring system that converts the rotational motion of the cams into linear motion to drive the linkages.

### Component configuration

The selection of an optimal number of cams and linkages is important to consider alongside the arrangement of internal parts and components, since Poka has space limitations. In our initial design, Figure 2(a), we considered two cams, each with two linkages. The cams were adjacent to one another and driven by separate motors. The initial position of the motors was offset by 180 °, resulting in an alternating tripod gait similar to that of DASH ^3^ and HAMR ^46^. This design would not fit inside the abdomen of the male specimen of *Eupatorus gracilicornis* due to the length of the cams. A modified version featuring only a single linkage per cam, Figure 2(b) was designed to reduce the size of the mechanism. However, the design was still too large to enable the installation of any further components and additionally, due to small differences in KV rating, the motors rotated at slightly different speeds, leading to desynchronisation of the legs. This caused the beetle to alternate between an alternating tripod gait when the offset of the legs was large, and a gallop when the legs moved closer together. In the second mode, no forward motion occurs, leading to inconsistent locomotion. To circumvent these issues a final configuration comprising only one linkage and one cam was selected, Figure 2(c), which had the benefit of extending the volume of available space inside the beetle for additional components, while eliminating the problems of desynchronisation as noted in prior configurations. This final design configuration is computationally rendered within the body of the beetle in Figure 3. The cam-linkage coupling in this final design is driven by one motor to enable walking. It also has a wheel driven by a second motor to enable turning. The principle of locomotion is similar to that of a walking dragline excavator ^58^. The motion of the mechanism can be divided into two phases: the stance and liftoff phases, Figure 4. During the stance phase, the foot of the mechanism is in contact with the ground, and it moves the robot forward. When the foot reaches the limit of its range of motion, it enters the liftoff phase, lifts off and the body makes contact with the ground. The foot moves forward, descends and engages with the ground, initiating the stance phase again and completing the cycle. This mode of locomotion results in a decreased energy cost. This is because only one motor is driven during walking, compared to two motors in previously described mechanisms. Furthermore, during the liftoff phase, the end effector is not in contact with the ground. As such the mechanism does not support the weight of the robot at this stage, and the motor does not have to counteract the weight. This results in a further decrease in potential energy consumption. A drawback of the mechanism is that it produces intermittent motion, as the robot only moves during the stance phase. This was deemed acceptable, as there were evident benefits in other aspects such as the reduction in size, of friction losses and of energy consumption. We used a foot mechanism with lateral legged reticulations rather than a separate-legged mechanism for two reasons. The first is that a foot mechanism allows us to minimise the internal components that required fitting within the space constraints of the beetle, since we only need to move a foot. The second is that the foot mechanism is structurally stable, and decreases the chances of catastrophic leg failure during payload bearing.

**Figure 2:**
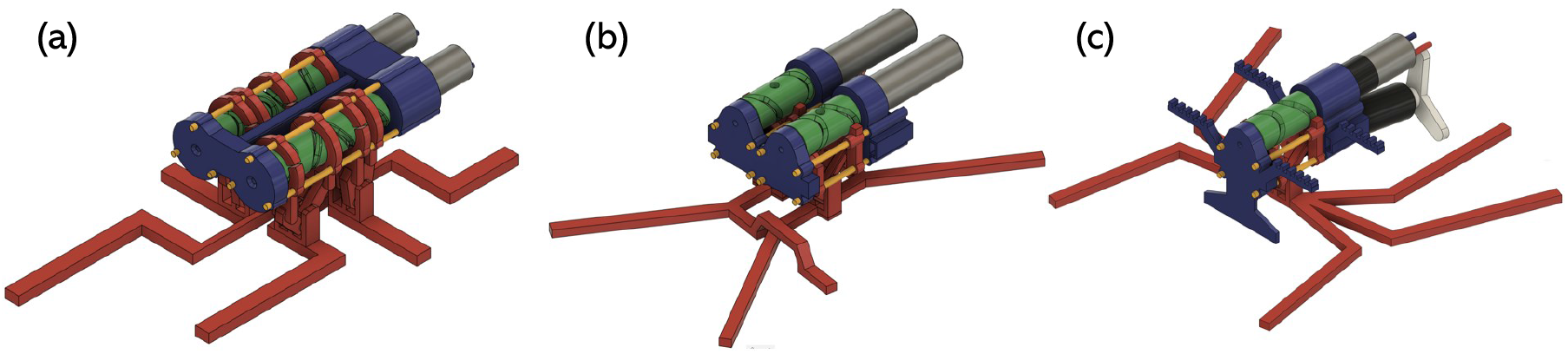
Design iterations of the necro-robot mechanism (to scale): (a) 2-cam (b) 2 linkages per cam; 2 cams, 1 linkage per cam; and (c) final mechanism with 1 linkage and 1 cam.

**Figure 3:**
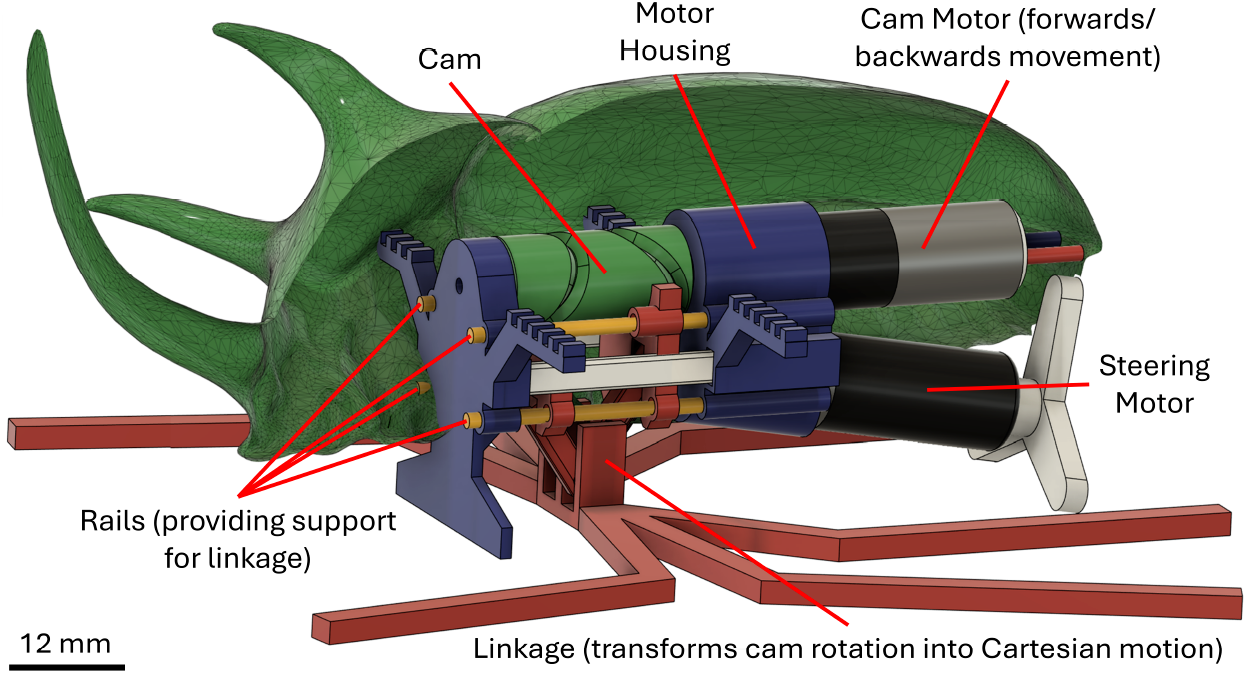
CAD render of the final ‘one cam one linkage’ design including all additional components.

**Figure 4:**
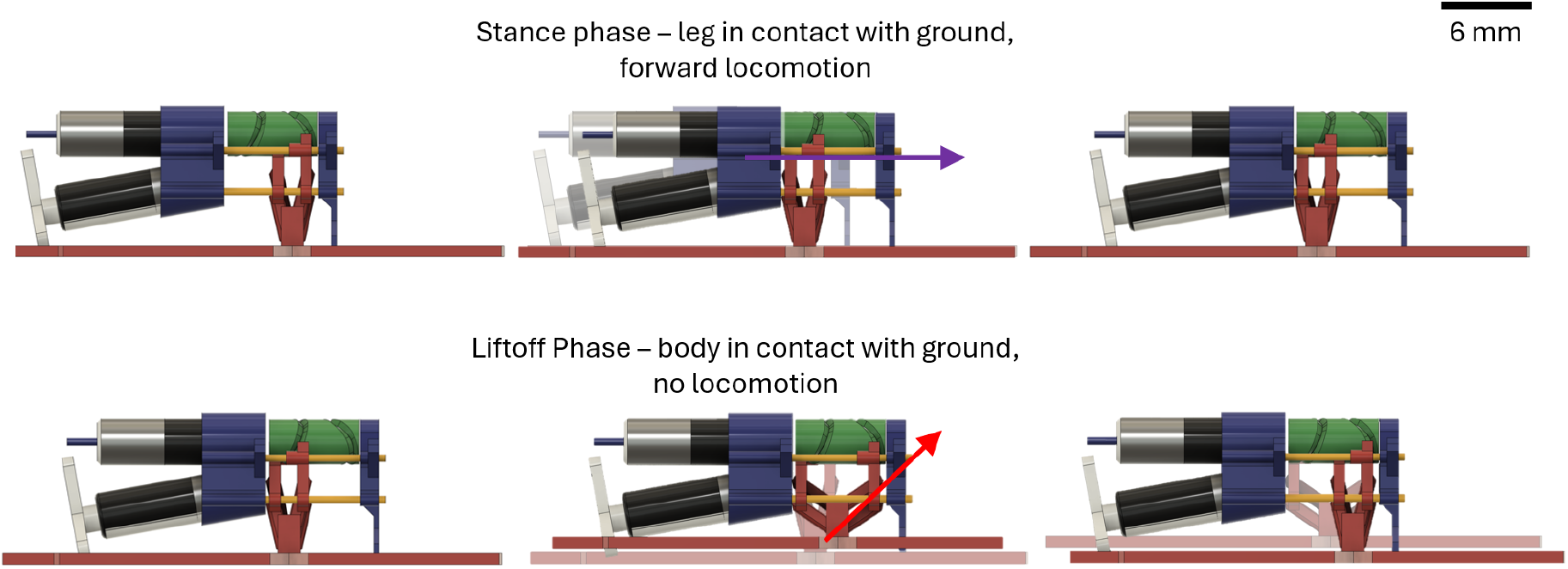
Timelapse visualisation of the stride phase (top) and liftoff phase (bottom) of the necro-robot gait.

**Figure 5:**
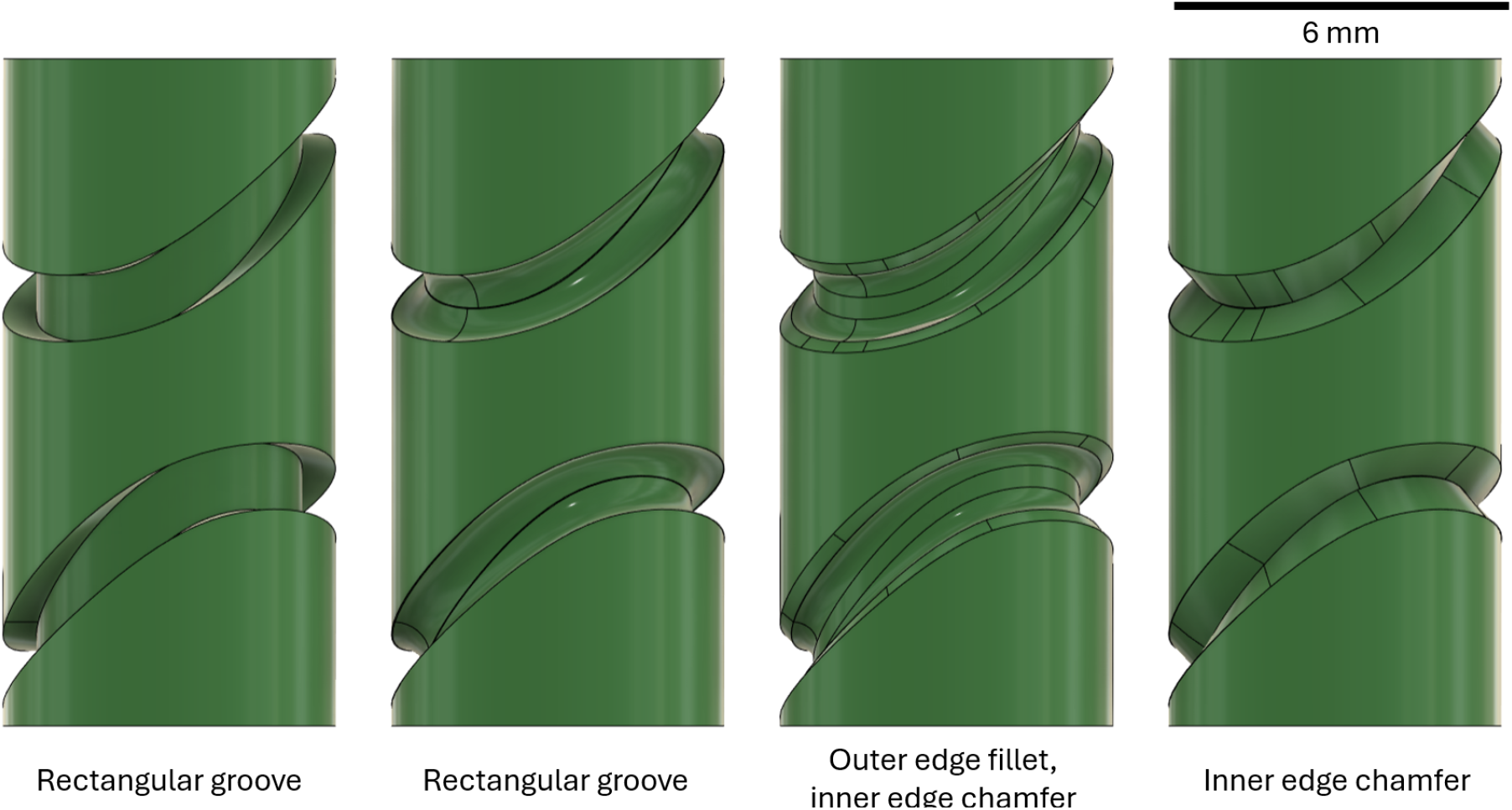
Cam variations explored to determine a feasible geometry for the groove.

**Figure 6:**
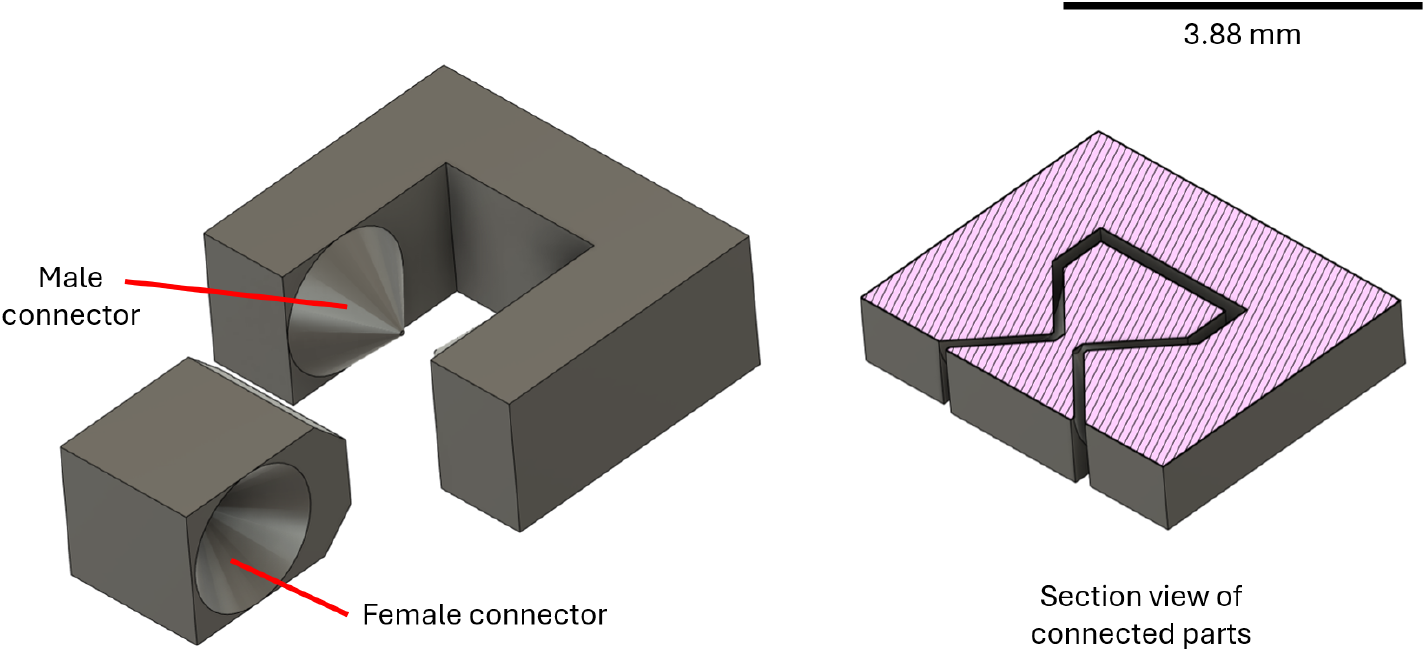
Print-in-place joint. Note the clearance between the parts in the section view.

### Cam design

The cam, is the machine component that controls the trajectory of the linkage through its interaction with the rings and is as such, a critical part of the actuation mechanism. Accurate manufacturing is therefore important and to prevent staircase effects from distorting the part, we printed the cylinder axis parallel to the *z*-axis. As such, the overhangs of grooves normally requiring supporting structures needed to be printed as accurately as possible. Three cam groove geometry solutions were assessed to improve print quality: (1) filleting of the inner grooves (2) filleting of the grooves in addition to the chamfering of their edges, and (3) chamfering of the grooves. The grooves were tested with a ring with a 1 mm conical pin. The cam was connected to the 2-stage gearbox motor, which was driven from an Arduino micro-controller. The motor was initially driven using 3.3 V, which was lowered until the motor was unable to drive the cam due to friction. We found that the ring could not be driven by either the filleted or filleted-chamfered solutions, while it was successfully driven by the chamfered cam. A cam with a diameter that is as large as possible for structural stability was designed to be balanced with its size, the cam itself never exceeding the overall dimensions of the mechanism. The cam diameter was 6 mm, matching the dimensions of the motor, as the mechanism as a whole must have a diameter at least as large as the motor to support it. The groove was manufactured to a depth of 0.5 mm and a width of 1 mm, to ensure that it could accommodate two 45° chamfers, effectively removing the distortion caused by the unsupported geometry.

### Linkage design

The creation of a complex linkage that can resist loads over 30× the weight of the robot while being feasible for 3D printing and possessing low friction posed a significant and novel design challenge. The only other milimetre-scale walking robots accomplish this through the smart composite manufacturing (SCM) procedure, which requires the use of a specialised laser with an ability to micromachine parts with typically, 1-10 *µ*m tolerances ^59^. These tolerances are orders of magnitude smaller than those from fused deposition modelling (FDM) 3D-printing, which can feature deviations within the range of 30-500 *µm*, depending on part orientation, feature size and printer calibration ^60^. Considering this, any linkage design would need to be printed with a distortion-resistant geometry. We consequently assessed two types of linkages: flexural linkages and rigid-body linkages. The initial flexural linkage designs consisted of a single flexure, featuring circular bending joints judiciously placed at desired degrees of freedom (DoF) locations. These linkages were unsuccessful since the flexures require sufficient bending forces, they also increased the torque requirement to actuate the mechanism. This resulted in significant reductions in efficiency. The forces required could not be decreased by thinning the flexures, as they were already manufactured from a single print layer, which is the minimal feature thickness that the printer could manufacture. In addition, the flexures were unable to locomote when loaded as they would deform, thus decreasing their range of motion.

Given the shortcomings of the flexure-based approach to the design of the linkages, rigid-body designs were investigated. Since the parts were too small to incorporate bearings or similar components to create low-tolerance revolute joints, we manufactured print-in-place 3D-printed joints. These consist of a male part with conical protrusions, which interact with similarly-shaped indentations in a female part. The joint was printed with its axis in the *xy*-plane, as the surface of the joint would otherwise be subject to the staircase effect, affecting surface smoothness from the limited resolution in the *z*-axis. To prevent the potential fusing of the parts, we included a clearance between the female and male parts. The exact value of this clearance depends on the 3D printer, which often varies between printers of the same model. As such, we deduced that it would be difficult to characterise the distortion analytically. To determine the clearances, multiple joints were tested iteratively with progressively increasing clearances. The final joint selected was the one with the smallest clearance that could be driven by the motor.

Early iterations of the print-in-place joint design were vulnerable to common defects in FDM 3D printing. Due to the ‘elephant’s foot’ defect, the bottommost layers of the part experience expansion due to insufficient cooling, or a build plate that is set too close to the nozzle. The expanded layers can lead to fusing on the print-in-place parts, or lead to joints being overly tight. To prevent this distortion from affecting the part, chamfers were applied to the bottom of the parts, extending up to 0.6mm (the first 5 layers of the print) above the surface in contact with the build plate, Figure 7. This effectively removed the effects of elephant’s foot defects and led to optimal clearances remaining consistent after build plate levelling. Due to the narrow and long geometry of the female parts, they were vulnerable to the manufacturing defect known as warping. As the part is being manufactured and the stacked layers cool, they contract, leading to edge peeling from the build plate. This can typically be resolved by enabling the printing of a brim, which consists of additional perimeters connected to the first layer of the part. Printing with a brim was deemed to be unfeasible, as it would increase considerably, the post-processing time of the parts, leading to imperfections at the edges that can in turn impede the joint movement. Instead, circular extensions to the bottom edges were added to the CAD models of the female parts. The brim was made circular, as any sharp edges could result in stress concentrations leading to easier detachment. We found a brim diameter of 5 mm was sufficient in preventing warping.

**Figure 7:**
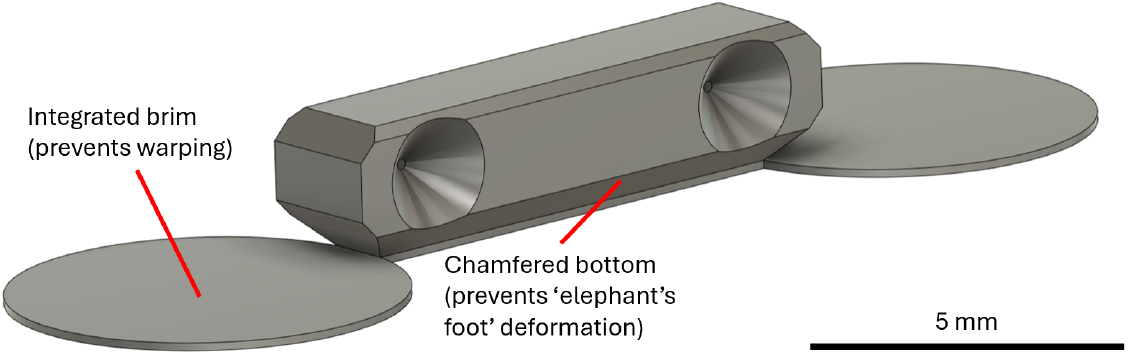
Linkage section with labelled integrated brim and chamfered bottom edge to prevent warping and elephant’s foot defects respectively.

### Ring system design

The rings are the elements that slide on the rails and interact with the groove of the cam. They are the main component responsible for transforming the rotary motion of the cam into linear motion to drive the linkage. Our initial iterations of the ring design considered 3D-printed rails to constrain motion linearly. This however proved to reduce efficiency due to the tribological properties of polylactic acid (PLA) used for printing. In plastic-on-plastic contact, polymers such as PLA experience static friction which is typically higher than kinetic friction. As the trajectory of the rings reaches point where there is null velocity due to closed-loop continuity, either increases in the required torque, or, discontinuous motion can result, reducing the overall smoothness of the ring as it follows the cam ^61^. Nevertheless, for plastic-on-steel contact, the static friction was noticeably lower than the kinetic friction ^62^, preventing the stick-slip motion from occurring. Steel-on-PLA contact also possesses a lower coefficient of friction than PLA-on-PLA contact, further reducing mechanical losses. These factors motivated our selection of 0.6 mm steel needles in the final design iteration for the rings, resulting in smoother overall motion. In the final design, Figure 8, the rings were only 1.5 mm thick, thus maximising their range of motion on the cam. The loose clearances needed to achieve low-friction motion potentialise problems of ring tilting on the rails, which may introduce positional errors of the end effector. To minimise this error, the length of one of the rail holes was increased to 3 mm, and an additional rail was introduced. Collision between the rings was prevented by isolating the rail hole on a rail that does not interact with the first ring. This improved stability without decreasing the range of motion generated by increasing the thickness of the rings. It was important to print the ring holes in a way that prevented material from entering. Initial versions featured these as holes integrated in an annular ring, which could not be printed. Observing G-code of this part, Figure 9, generated by the slicer revealed the cause to be the intersection of the line, which defined the hole with the perimeter of the part. The lines intersecting generated local material build resulting in a thicker layer line. This in turn decreased the diameter of the printed ring holes. To circumvent this, we resolved the problems of intersecting lines by generating a single line along the perimeter of the hole. To ensure there was no distortion caused by warping, brim-like extensions were added around these perimeters, also improving adhesion to the build plate. The geometry of the feature of the ring that interacts with the cam is of significant importance to the load-carrying capacity of the necro-robot. Initially, cylindrical and conical pins were used, which were slotted inside the cam groove. However, running them under load for short periods of time (≈5 min) led to pin wear, and ring slippage from the grooves, making locomotion impossible. We noted that the conical pins resulted in features shorter than the layer line width of 0.4 mm. As such, they were impossible to manufacture in full. To circumvent these issues, a V-shaped pin was designed into the ring. The V-shape takes advantage of the fact that while the minimum feature length is 0.4 mm, the layer height is lower than that at 0.12 mm. This means that the tip of the V-shape can be successfully printed, as long as the edge is longer than 0.4 mm. The V-shaped pin is also resistant to wear as noted during experimentation.

**Figure 8:**
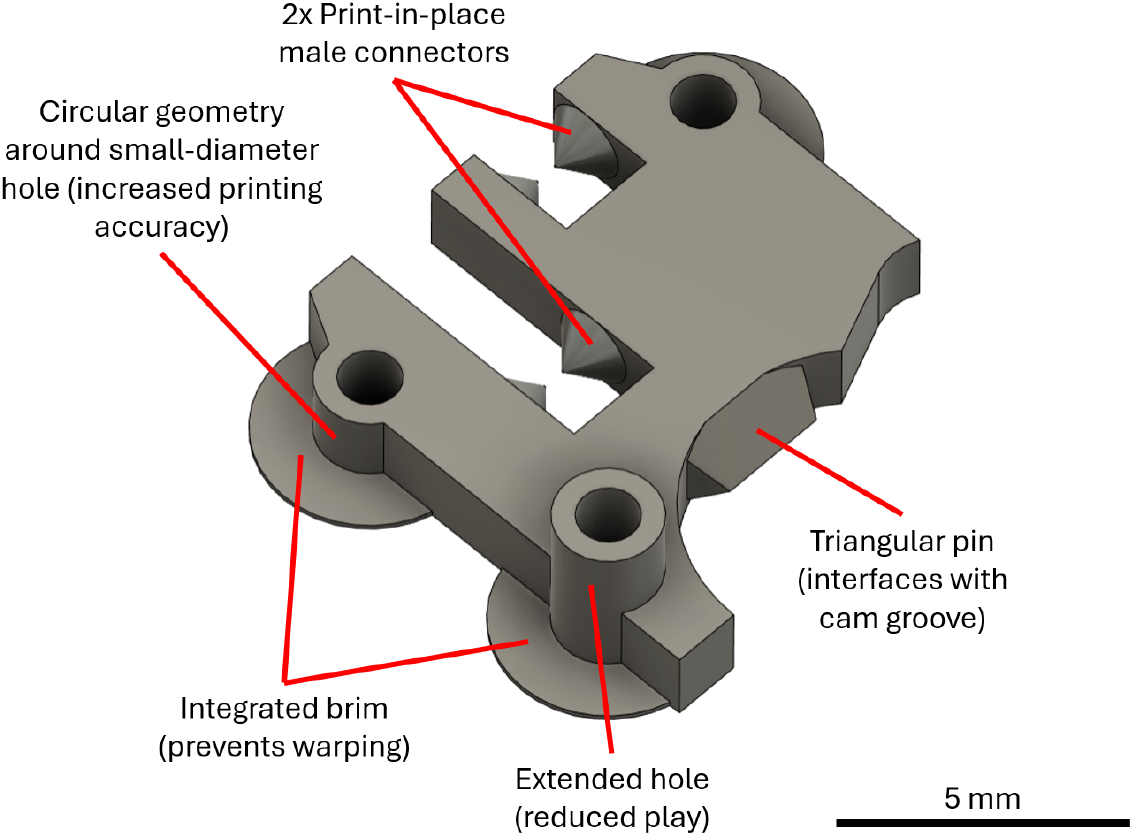
Final ring design with labelled features of interest (clockwise): print-in-place male connectors, triangular pin for cam engagement, extended rail hole, integrated brim, circular rail hole wall.

**Figure 9:**
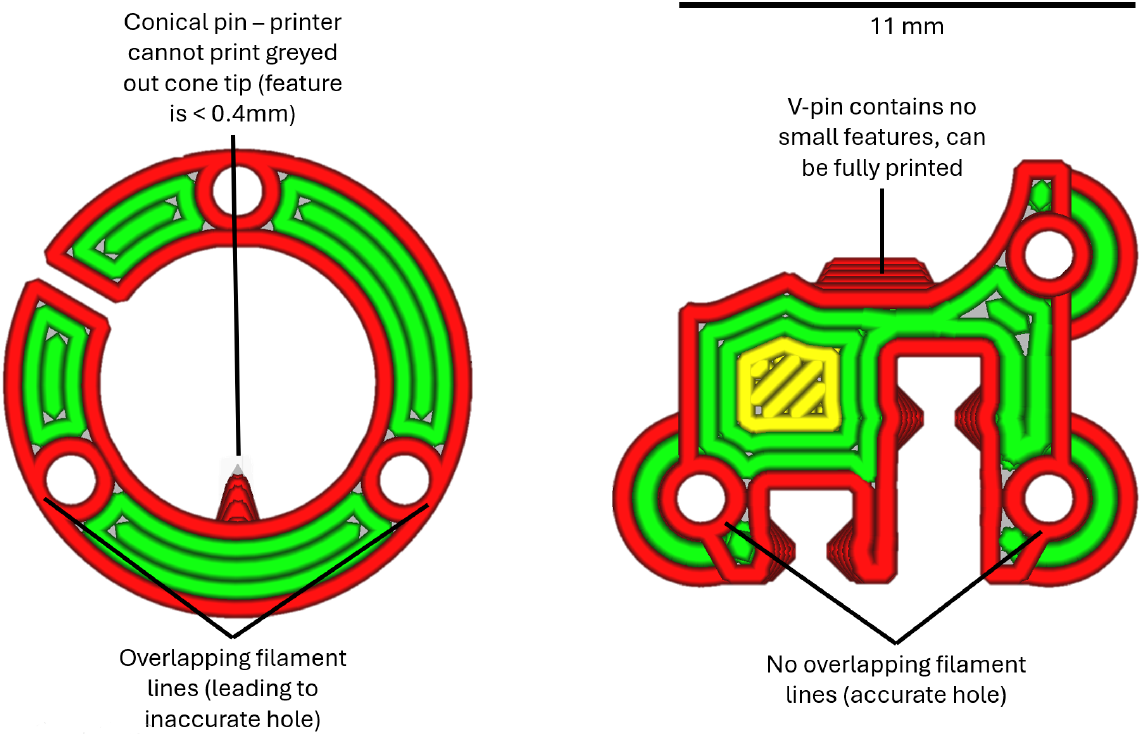
Comparison of the G-code for the final ring design with an older ring. A reduction of distortion is achieved through the use of G-code compatible geometry.

### Actuator selection and motor assembly design

The selection of appropriate lightweight actuation is an important step towards an optimised payload ratio. The actuator requires high torque yet should be small enough to fit inside the beetle together with the ambulation mechanism. At the millimeter length scales, DC motors and piezoelectric actuators have already demonstrated potential. Piezoelectric actuators, such as those used by HAMR, would be unsuitable as they require high-precision manufacturing methods such as SCM. We selected 6 mm diameter drone DC motors, which come with reducers that are required to supply sufficient torque for the movement of heavy loads. Two different reducer-motor packages were considered, comprising either a 2-stage (2×) and a 4-stage (4×) planetary gearbox. Table 2 summarises their properties.

**Table 2:** Specifications of the motors with 2*×* and 4*×* gearboxes, respectively.

|  | Nominal voltage [V] | Speed at 3V [rad/s] | Torque [g-cm] |
| --- | --- | --- | --- |
| 2x gearbox | 3.3 | 126 | 27 |
| 4x gearbox | 3.3 | 4.81 | 180 |

The motor assembly (motor housing, rails and end cap), bears the load of the mechanism which is integrated with the body of the beetle. The motor assembly should therefore comprise features to enable direct contact with the shell. To achieve this, toothed extensions were manufactured onto the motor housing and ring. When cut to the appropriate size, this allows the beetle to act as a support for the mechanism and the motor housing was attached to the inside of the beetle using cyanoacrylate glue. This set up turns the body of the beetle into a load-bearing structure, thus taking advantage of the high stiffness of the hardened parts of the beetle body (sclerites).

### Trajectory mapping

For each motor, two sets of eight experiments were performed, one with randomised but non-repeating linkage-cam pairs (samples 1-8), and another where all cams were tested using the same linkage (samples 9-16). Figure 10 provides as an example, a sequence from trajectory analysis filming, where one elliptical cycle was completed. In both sample sets, the cams were tested in the same order, such that the cam from sample 1 was used again in sample 9. Each trajectory consisted of 600 points, representing at least 10 revolutions. The error was evaluated as the minimal distance between a given experimental point and any point of the theoretical ellipse for 2× and 4× gearbox motors, respectively. On average, the experimental trajectory deviates by 0.087 mm, which represents ca. 6% of the minor diameter and 3% of the major diameter. It is of note that the 2× gearbox motor trajectories, Figure 11(a), possess a slightly higher mean error than the 4× gearbox motor trajectories, Figure 11(b). This, combined with the extended interquartile range observed for the 2× gearbox motor, suggest that the dynamic behaviour of the linkage leads to a minor decrease in accuracy. The mean errors for all sample sets are summarised in Table 3. There are two other significant sources of error: the tolerances of the manufacturing method, and distortions independent of the geometry of the part. The first is presumably responsible for the majority of trajectory errors in the system. An analysis of the dimensional accuracy of fused deposition modelling (FDM) 3D printing shows that consumer-grade 3D printers are able to manufacture the complex curves associated with conventional cams, to an accuracy of ± 0.06mm in the *xy*-plane ^63^, which accounts for the majority of the error observed. The second source of error (distortions independent of the geometry part) could account for the deformations of the trajectories, which are consistent across both the 4× and 2× motor experiments, such as the particular trajectory shape exhibited for sample 4 in both sample sets. Since the cams were manufactured in a single batch, significant stringing was observed on the parts. Stringing is a manufacturing defect consisting of thin strings of filament caused by the nozzle material percolating during travel. This in turn typically results in the formation of globules of material within the grooves of the cam, leading to deformation of the mechanism as the linkage travels over them. A proof detailing the accuracy of the cam is provided (Electronic Supplementary Material: Proof of Cam Accuracy), where comparisons are made against other millimetre-scale robots employing 3D-printed cam mechanisms ^64^. Further, we provide detail on the interchangeability of the cam-linkage couplings (thus relative reliability), comparing randomly paired camlinkages against one specific linkage paired and tested with each individual cam (see Electronic Supplementary Material: Statistical Analysis on the Interchangeability of the Cam-Linkage Couplings).The internal grooves of the cam were lined with dry graphite to decrease friction at the interface of the cam-linkage coupling.

**Table 3:** Summary of the mean errors for all sample sets for each of the motors. The first row of data represents samples 1-8, while the second row represents samples 9-16.

|  | 4x gearbox | 2x gearbox |
| --- | --- | --- |
| Mean error, randomised cam and linkage [mm] | 0.0821 | 0.1083 |
| Mean error, randomised cam only [mm] | 0.0727 | 0.0885 |
| Mean error overall [mm] | 0.0773 | 0.0964 |

**Figure 10:**
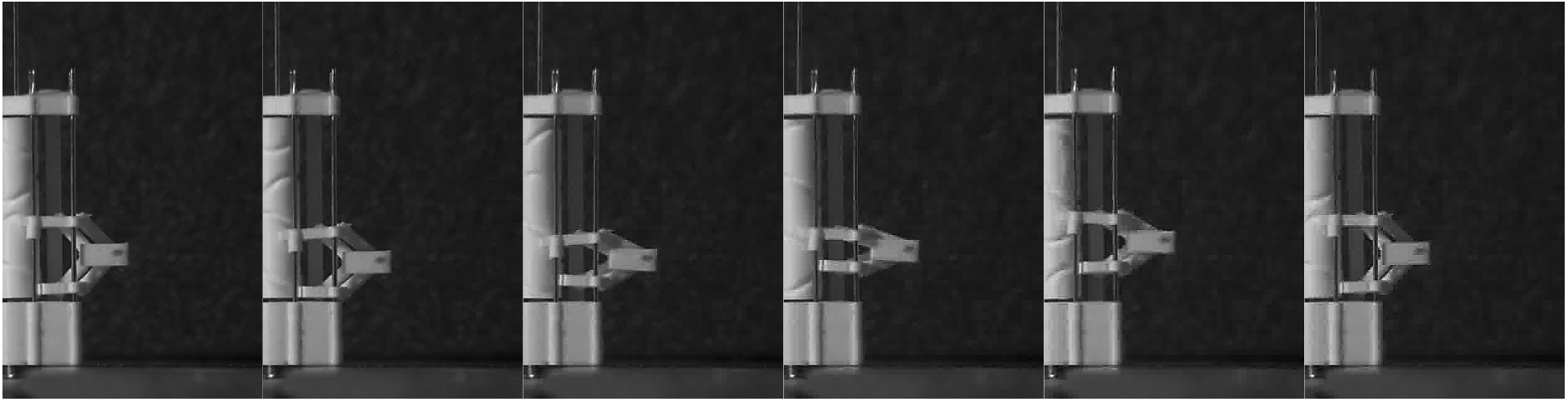
Sequence of images from a single motion cycle (for a 2 *×* ear) during trajectory filming. The full sequence of can be viewed (see Electronic Supplementary Video 1).

**Figure 11:**
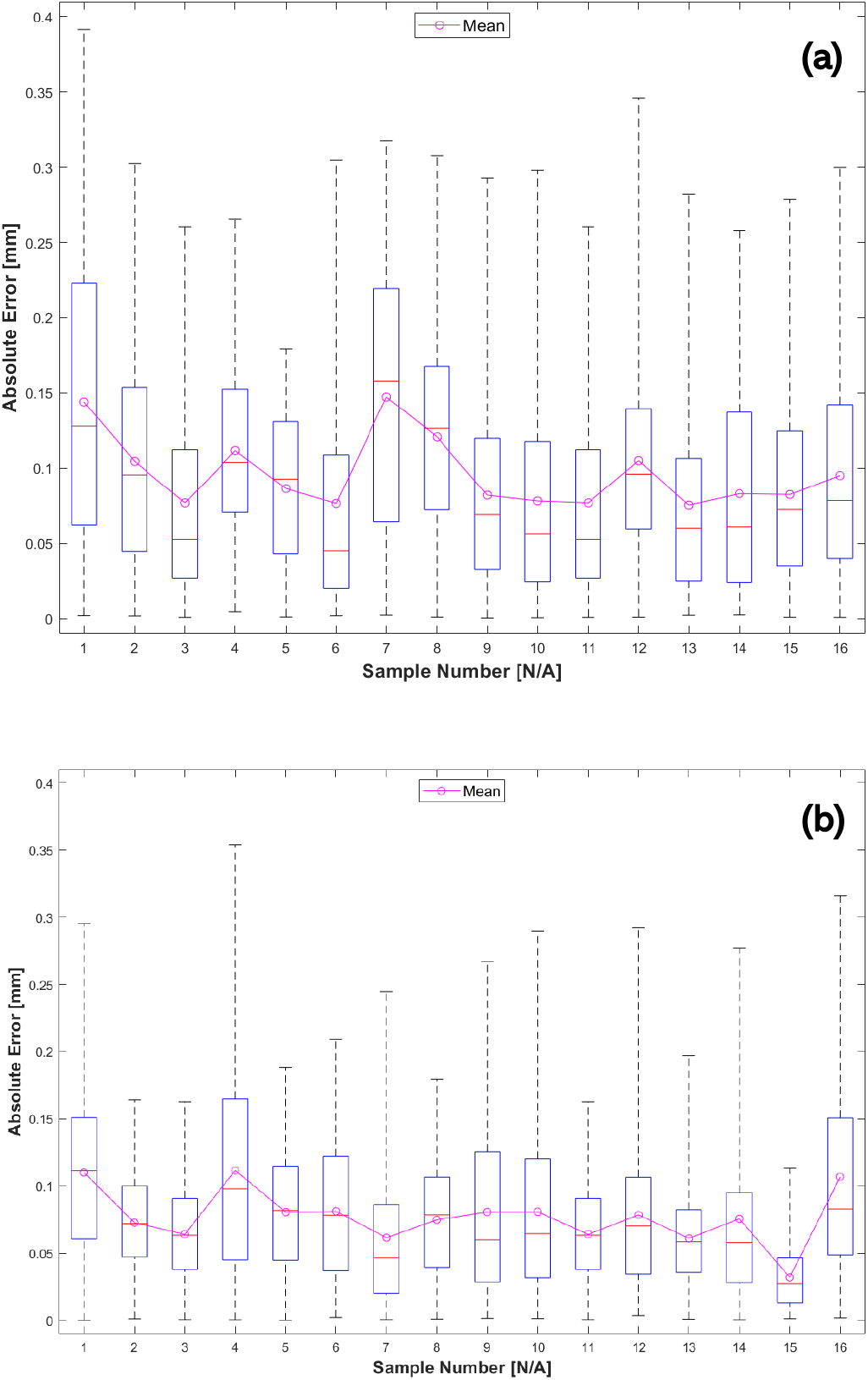
Box and whisker plot of the trajectory error across all samples for (a) the 2*×* gearbox motor and (b) the 4*times* gearbox motor. The error represents the shortest distances between any point and the theoretical ellipse. The red line in each box plot defines the median value, and the whiskers denote the extremes of the set. The heights of the boxes represent the interquartile range,within which 50% of the data falls. The individual trajectories mapped against the ellipses for both 2*×* and 4*×* motors, are provided (see Electronic Supplementary Material: Individual Trajectories Mapped against the Ellipses for both 2*×* and 4*×* Motors).

### Payload experimentation

The water in the bottle was increased in increments of 100 g, starting with a payload weight of 200 g. Eight trials (n = 8) were conducted at each payload weight to verify consistency. A sequence of Poka carrying a 500 g payload weight is shown in Figure 12. We obtained 250 samples from the middle of each data set to capture the steady-state current consumption, corresponding to at least eight full revolutions of the cam. The data from each payload weight was then concatenated to preserve the maximal and minimal values, since averaging would reduce the extrema. The necro-robot Poka was able to carry a maximum payload of 500 g, equivalent to 6847% of its body weight. We note there is a linear relationship between payload weight and current consumption, with a coefficient of determination *R*^2^ = 0.9856, Figure 13.

**Figure 12:**
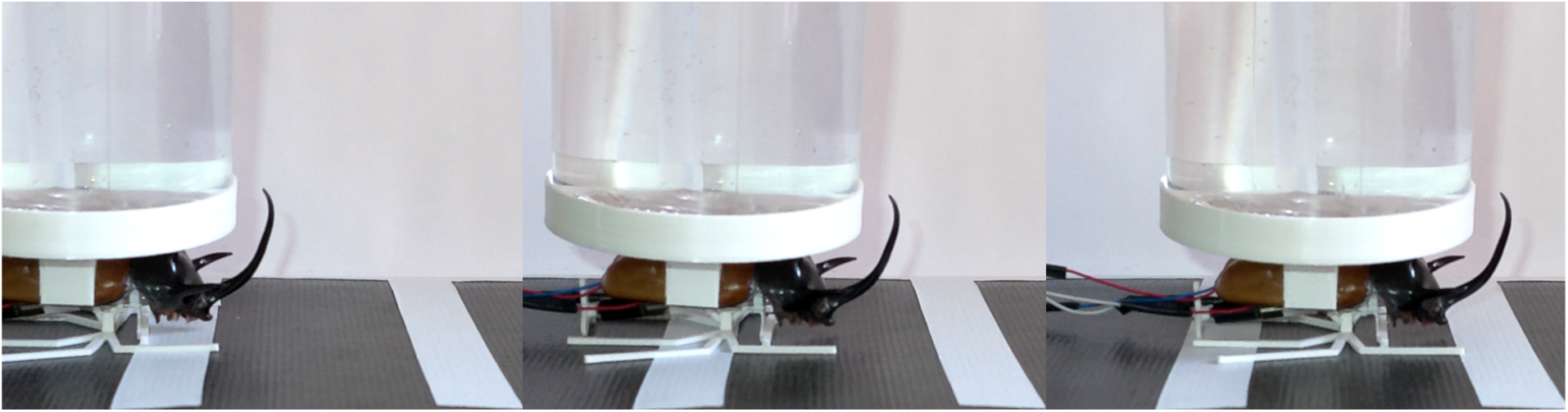
Sequence of our necro-robot Poka carrying a 500 g payload. To watch Poka carrying a 500g payload, refer to the Electronic Supplementary Video 2.

**Figure 13:**
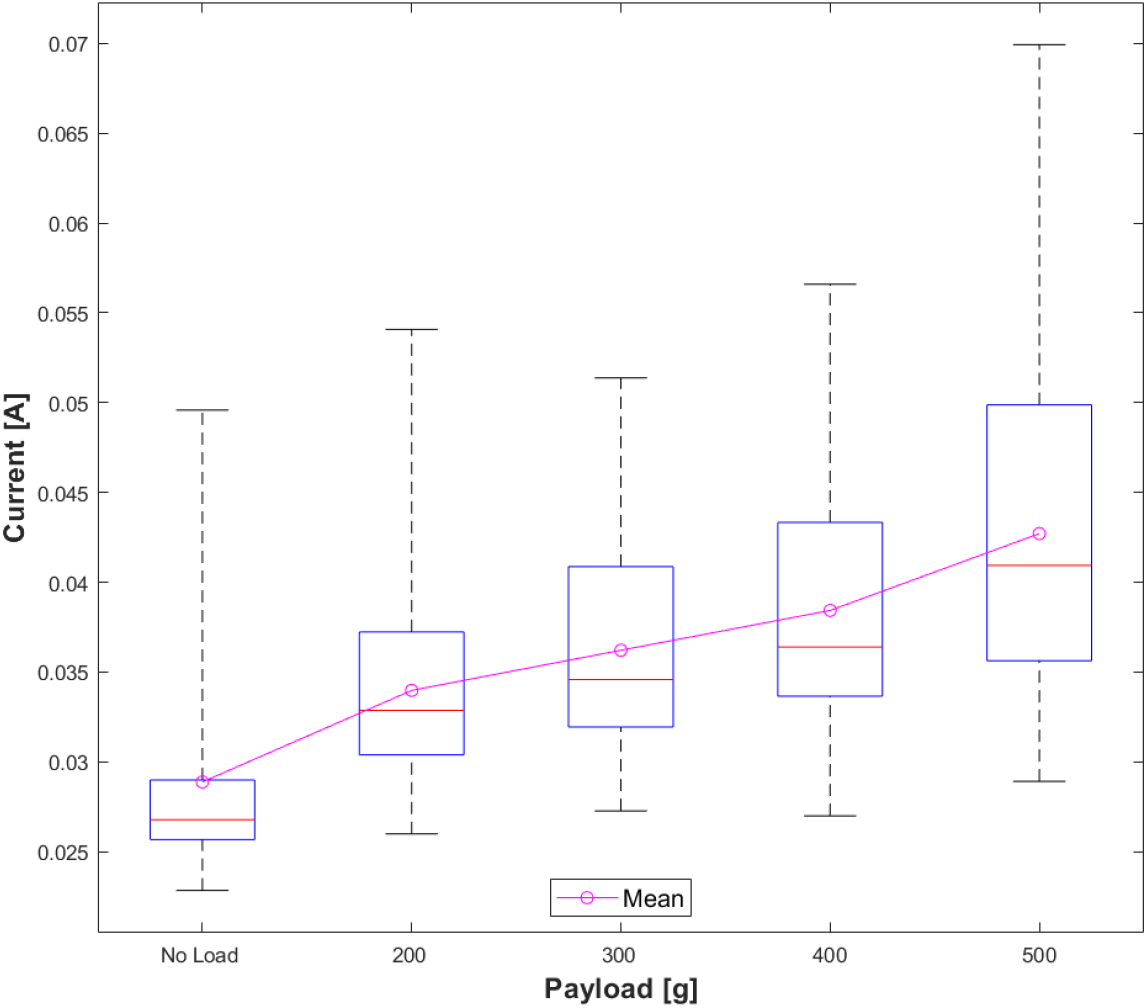
Box-whisker plot of the current consumption vs. its payload. The red line in each box plot denotes the median value, and the whiskers denote the extremes of the set. The heights of the boxes represent the interquartile range - the range in which 50% of the data falls. The current data increases linearly with payload, with *R*^2^ = 0.9856 (cf. Electronic Supplementary: Current Data Increasing Linearly with Payload).

Poka’s average walking speeds were determined at different payloads, Figure 14, between two markers spaced at a distance of 5 cm apart. At a 500 g payload, Poka’s speed is reduced to approximately 87% of its unloaded speed. It is interesting to note that Poka’s speed was higher when carrying a payload of 200 g (27.4 times its body weight), than it was in its unloaded state. This suggested that payload improves Poka’s ability to gain greater surface traction, and that in the absence of payload, Poka slips rather than grips.

**Figure 14:**
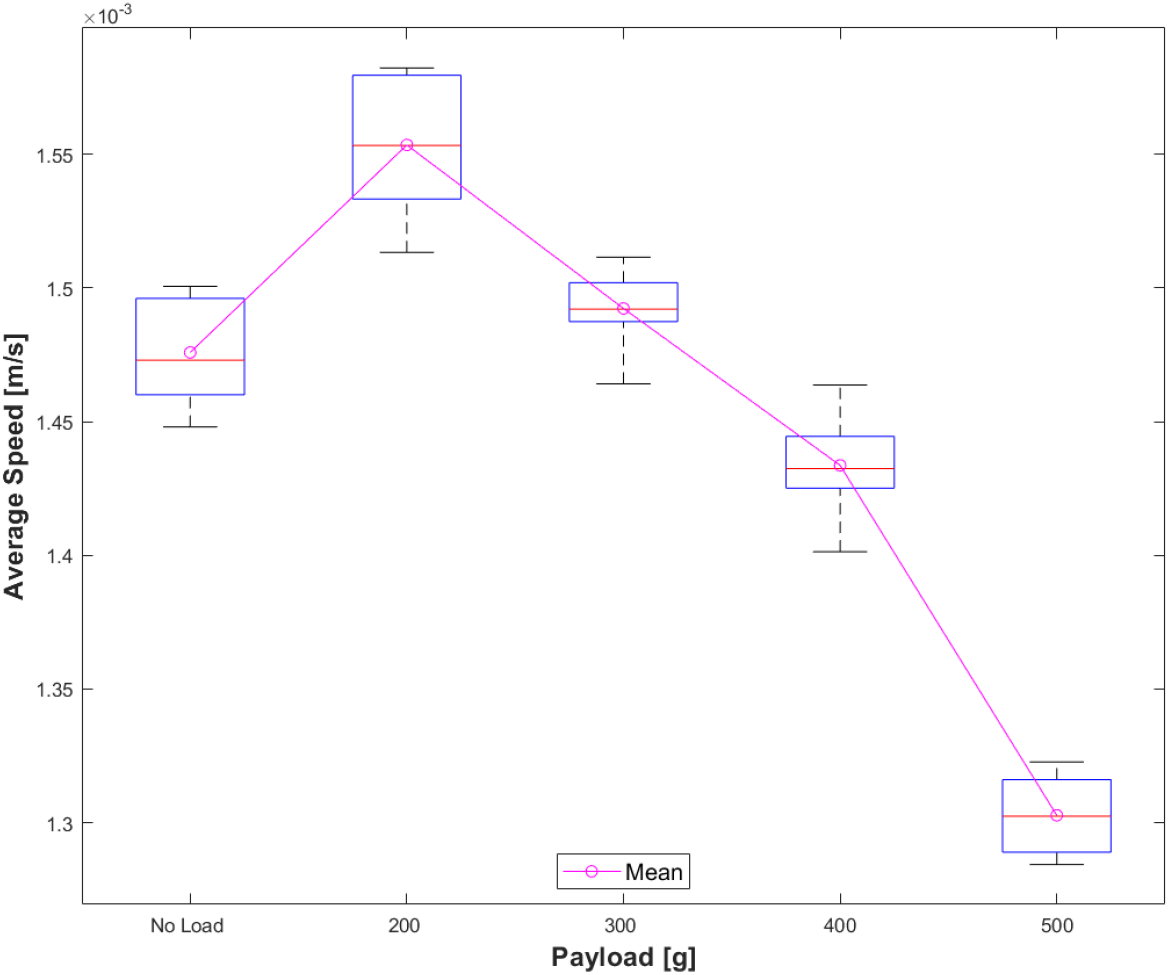
Box-whisker plot of the necro-robot speed vs. its payload. The red line in each box plot denotes the median value, and the whiskers denote the extremes of the set. The heights of the boxes represent the interquartile range - the range in which 50% of the data falls.

When comparing against real rhinoceros beetles, using the data collected by Kram and co-workers ^38^, Table 4, the necro-robot Poka significantly outperforms *Xyloryctes thestalus* in terms of maximum payload carried, *m*_*p*_, and the consequent payload ratio, *PR*, which is a ratio of the maximum payload to the body mass 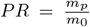. While Poka’s body mass is on average 307% above that of *Xyloryctes thestalus*, its maximum payload carrying capacity is 700% higher, and its consequent payload ratio is 228% greater. Nevertheless, the rhinoceros beetle is able to carry its payload at a significantly lower specific power, 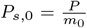, ca. 3-fold less power per unit body mass. However, when normalised against the total mass, the specific power of Poka, 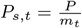 (where *m*_*t*_ = *m*_0_ + *m*_*p*_), is of the same order of magnitude as that of the beetle *Xyloryctes thestalus*, being only 33% higher in Poka.

**Table 4:** Load carrying performance comparison between the European rhinoceros beetle (*Xyloryctes thestalus*) and our necro-robot, Poka. The data for the rhinoceros beetle is sourced from Kram et al. ^38^.

|  | *Xyloryctes thestalus* | Necro-robot, Poka |
| --- | --- | --- |
| Body mass, $m_0$ [g] | 2.38 | 7.302 |
| Maximum payload, $m_p$ [g] | 71.4 | 500 |
| Payload ratio, $PR$ [%] | 3000 | 6847 |
| Specific Power (power per unit body mass) at max payload, $P_{s,0}$ [W/kg] | 6.48 | 19.24 |
| Specific Power (power per unit total mass) at max payload, $P_{s,t}$ [W/kg] | 0.21 | 0.28 |
| Power per unit body mass at no payload [W/kg] | 1.66 | 12.89 |

There is also relevance in comparing Poka against other ambulating robots. It is important to have an ability to traverse terrain in an efficient manner, especially for longer, autonomous operations. As the necro-robot Poka can be used in such a manner, we will first compare its energy efficiency. The metric chosen for this comparison is the cost of transport (CoT), Equation (1), which is a popular scale-agnostic metric used in the determination of gait efficiency, where 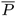 is the average power consumption, *m* is the mass of the robot and 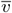 is its average velocity for the power consumption. *P* is calculated by multiplying the mean current consumption measured in the payload experiment with the 3.3*V* voltage used in the experiments. Table 5 provides the *CoT* values for Poka at different payload weights.

**Table 5:**
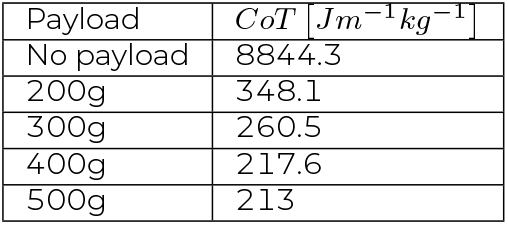
Cost of transport, *CoT*, calculations for our necro-robot Poka under different payloads.

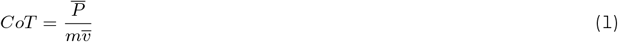

Comparing the two extrema of the data against other extreme mobility robots, Table 6, reveals that when loaded, the necro-robot Poka has a significantly lower specific power 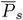 than any other ambulating robot. In contrast, the *CoT* is orders of magnitude higher than that of the other robots in both the loaded and unloaded cases. This, combined with the low 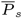, suggests that the cause for the large *CoT* value, is the low speed of Poka. Slow locomotion in Poka is a function of the relatively weak motor used (as judged by the power consumption), and the small radius of the cam, limiting the maximal slope of the cam groove and thus the actuation speed of the mechanism. Nevertheless, where comparing the payload ratio, *PR*, of Poka against other ambulating robots, Figure 15, we find that Poka’s *PR* at 6847%, far exceeds the highest of any other robot reported to date, to the best of our knowledge, with the subsequent highest *PR* belonging to Superbot ^65^, who has a *PR* of 530%, a factor of 13 lower than that of Poka. This finding highlights the benefits in (a) the use of low density materials (such as PLA) to manufacture bionic parts and components into the necro-robot, and (b) the utility of insect exoskeletons, which are low density chitin-reinforced composites with exceptional specific properties ^66–68^ and lightweight architectures ^69–72^. We additionally note from Figure 15, a non-linear relationship between *PR* and *m*, as evidenced by the Spearman’s coefficient of -0.63 and p ≤ 0.01. The determination coefficient of 0.46 indicates there is a high level of scatter about the regression fit, which is of the form *PR* = e^*a* ln(*m*)+*b*^, where for the current data set used in this figure, *a* = -0.34 and *b* = 0.35.

**Table 6:** Comparison of 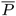, *m, v, CoT* and 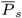 for the necrobot Poka under unloaded and maximum payload, DASH, Mini-Whegs, RHex and iSprawl robots.

|  | Necrobot | Loaded Necrobot | DASH<sup>3</sup> | Mini-Whegs<sup>73</sup> | RHex<sup>74</sup> | iSprawl<sup>75</sup> |
| --- | --- | --- | --- | --- | --- | --- |
| $\bar{P}$ [W] | 0.094 | 0.14 | 0.36 | 1.2 | 80 | 12 |
| $m$ [g] | 7.3 | 507.3 | 16.2 | 146 | 7500 | 315 |
| $v$ [m/s] | 0.001475 | 0.0013 | 1.5 | 0.9 | 0.5 | 2.3 |
| $CoT$ [Jm<sup>-1</sup>kg<sup>-1</sup>] | 8844 | 213 | 14.7 | 9.132 | 21.3 | 17.4 |
| $P_s = P/m$ [Wkg<sup>-1</sup>] | 12.87 | 0.28 | 22.0 | 8.57 | 10.7 | 38.1 |

**Figure 15:**
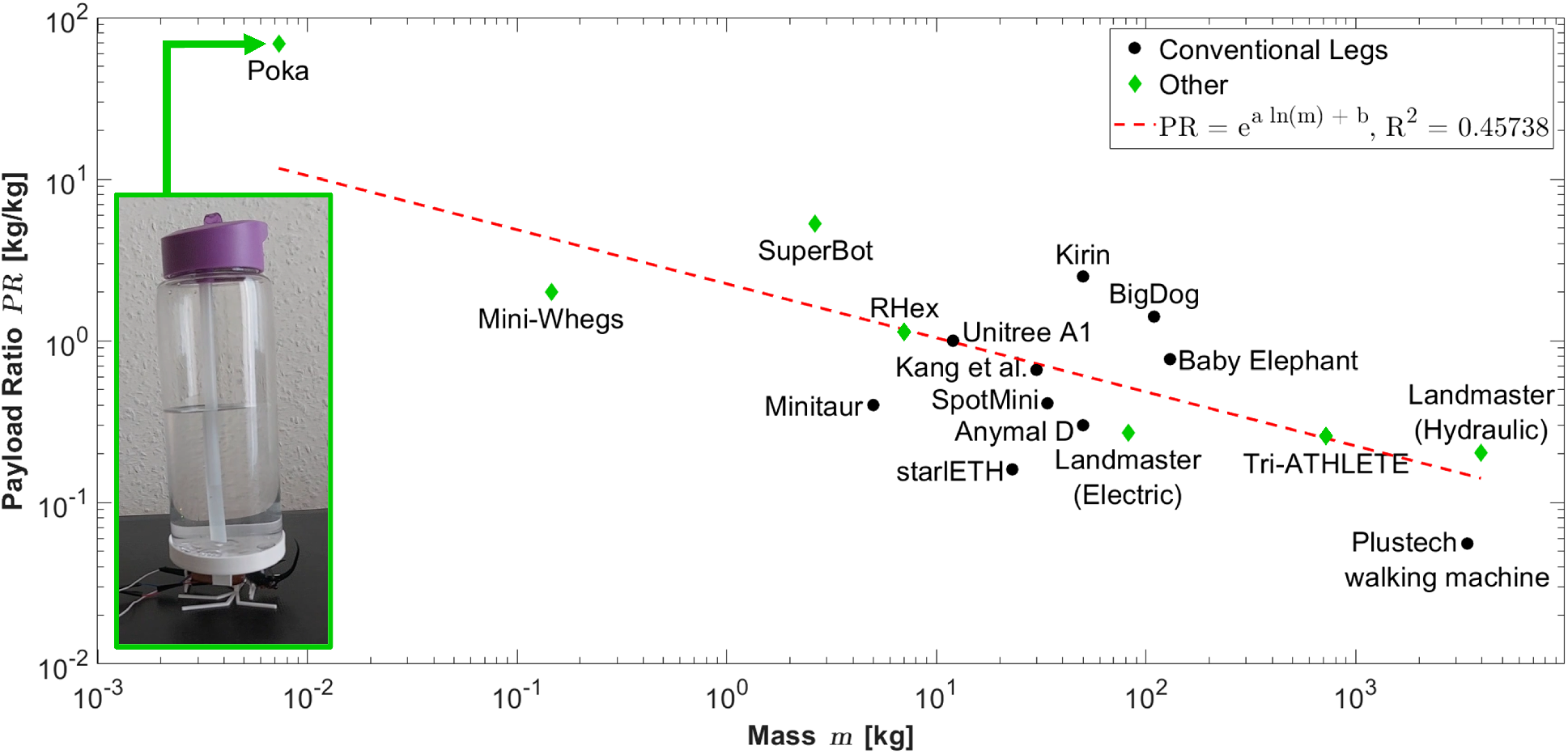
Payload ratio, *PR*, plotted against robot mass, *m*, for a broad variety of robots, showing also the position of Poka relative to all other robots. The data for robots other than Poka in this plot is either taken directly from, or deduced from data available in the following papers: ^65,73,74,76–87^. For the dataset elucidated in this plot, *PR* is shown to be non-linearly related to *m* as: *PR* = e^*a* ln(*m*)+*b*^ where *a* = -0.34 and *b* = 0.35. A table providing numerical values used in this graph alongside their sources, is provided (cf. Electronic Supplementary Material: Tabulated Payload Ratios of Poka Compared against Numerous Robots).

## CONCLUSIONS

Our original aim was to design, manufacture and validate a necro-robot beetle with an extreme payload ratio, *PR*, matching that of a live rhinoceros beetle, *Xyloryctes thestalus* at *>* 3000%. We have far exceeded this by reaching a measured *PR* = 6847%, meaning that our beetle necro-robot (dubbed ‘Poka’) is able to ambulate while carrying ca. 68.5 × its own body weight. We use the exoskeleton of a deceased five-horned rhinoceros beetle (*Eupatorus gracilicornis*) as a chassis, inside of which we mechatronically engineer an actuation mechanism comprising a single cam, a single linkage, a single motor and additional supporting components. While our *PR* is significantly higher than that of a live rhinoceros beetle, *Xyloryctes thestalus*, the specific power at maximum payload, *P*_*s,t*_, is of the same order of magnitude between the a real beetle (0.21 W/kg) and Poka (0.28 W/kg), indicating that the power efficiency of Poka per unit weight is similar to that of a real beetle, even though its *PR* is more than 2-fold superior. Poka’s highest average speed, 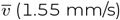 is achieved at a *PR* = 2739%, after which it progressively decreases with increasing payload ratio, reaching its minimum 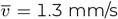 at maximum payload ratio. We furthermore compare the *PR* of Poka against sixteen other ambulating robots and find that when plotted against the mass of the robot, a non-linear relationship can be mapped between the two following the form *PR* = e^*a* ln(*m*)+*b*^, where for the data set plotted, *a* = -0.34 and *b* = 0.35. The data yields a Spearman’s coefficient of -0.63 and a p ≤ 0.01, indicating negative correlation between the two variables, while a coefficient of determination is calculated at 0.46, indicating there is still high scatter about the regression line. Of all the robots plotted for *PR* against mass, Poka is shown to have the lowest mass and the highest *PR* (6847%), exceeding even that of SuperBot (530%), whilst also having the lowest mass of any of the robots. To the best of our knowledge, the necro-robot beetle ‘Poka’, has by far, the highest recorded payload ratio to date. We consider this is related to (a) the use of the beetle exoskeleton as a chassis, as it is a natural composite with high specific properties, and (b) the use of stiff polymer, geometrically optimised and additively manufactured into structurally stable parts for the necro-robot. Our outputs provide strong justification for continued research into the repurposing of insect bodies to create millimetre-scale robots with high performance and low cost.

## MATERIALS AND METHODS

### Beetle procurement, conditioning and initial morphological surveys

Dessicated five horned rhinoceros beetles (*Eupatorus gracilicornis*) were purchased from Bugs Direct Ltd (Buckfastleigh, UK; Company number: 06181289). Specimens were relaxed in a humid environment using a moisture chamber ^88^ for three days. Moisture softens intersegmental membranes and enable the limbs to be moved. This in turn enabled a preliminary check of joint flexibility, as well as beetle dissection, allowing us to survey possible integration locations for electronic and mechanical components within the beetle. The abdomen was dimensionally surveyed as it contains the largest internal volume that would be able to house electro-mechanical components, while the beetle elytra are rigid structures that could protect the components if inserted into the abdominal cavity. A small free internal volume in the pronotum was surveyed, as this could be increased by removing internal matter. The pronotum is also protected by the thickest chitin layer found on the beetle, potentialising its utility as a high load bearing structure. The legs could not be actuated without breakage and they were deemed too brittle to include in the necrorobot and were thus excluded from further evaluation. The positions of the protonum and elytra are provided (cf. Electronic Supplementary Material: Separated Body Parts of the Five Horned Rhinocerous Beetle).

### 3D scanning and computer aided design (CAD)

We 3D scanned and computationally rendered the geometrical construction of the beetle to conceptually design the judicious fitting of an electro-mechanical package within the necro-robot. For this purpose, we removed matter from the abdomen and protonum, as it would allow us to conceptually evaluate dimensional limitations and opportunities. A portable 3D scanner (Einscan Pro 2X Plus) was first used in fixed scan mode, with the beetle mounted on a turntable. This mode was chosen since it offers the highest scanning accuracy of 0.04 mm ^89^. The beetle exoskeleton features dark reflective surfaces, which are particularly challenging to scan ^90^. This was resolved by applying talcum powder with a brush to the surfaces, resulting in an easy-to-scan matte surface.

### Materials, manufacture and component selection

Based on both the 3D scan of and from dissection, we deduced that the abdomen and pronotum offer sufficient internal volume to house a small DC motor (Garosa 6mm Micro Motor) and an actuation mechanism comprising linkages, cams and supporting components. The components were therefore selected based on both the functional payload requirements and the space constraints imposed by the internal structure of the beetle. Fused deposition modelling (FDM) 3D printing was selected as the manufacturing method. We computationally sliced our mechanism in Cura (Ultimaker B.V., Utrecht, Netherlands) and the mechanism was manufactured on a Kingroon KP3S (Shenzhen Kingroon Tech Co. Ltd., Kowloon, Hong Kong), with a layer height of 0.12mm. This is the lowest layer height for which the manufacturer provides a tuned slicing profile.

### Motion generation: theory

The ambulation of Poka was enabled by designing with respect to the trajectories of the end effectors. As the motion of the mechanism is driven fully by the cam, it follows that the motion of the robot should be defined by the geometry of the cam rather than through programmed actuators, driven in systematically, as is the case with conventional robot ambulation. Designing the cam grooves for a particular motion was achieved by defining the relationship between the Cartesian position of the end effector and the positions of the rings (i.e. the inverse kinematics), Figure 16. This was followed by the substitution of the equations of the desired curve into the inverse kinematics equations. Starting with a coordinate system with an origin at the leftmost position of the left ring, the double four-bar mechanism could be simplified to a trapezoid linkage with a middle element parallel to the *x*-axis. We were interested in finding the positions of the left and right rings, *x*_1_ and *x*_2_ respectively, expressed using the coordinates of the midpoint of the end effector, *x* and *y*. The length of the trapezoid legs is marked by *l* and the shorter base length by *k. x* then corresponds to the midpoint of the large base of the trapezoid and *y* is expressed using the Pythagorean theorem on the triangle ACO, Equation (2) and Equation (3).

**Figure 16:**
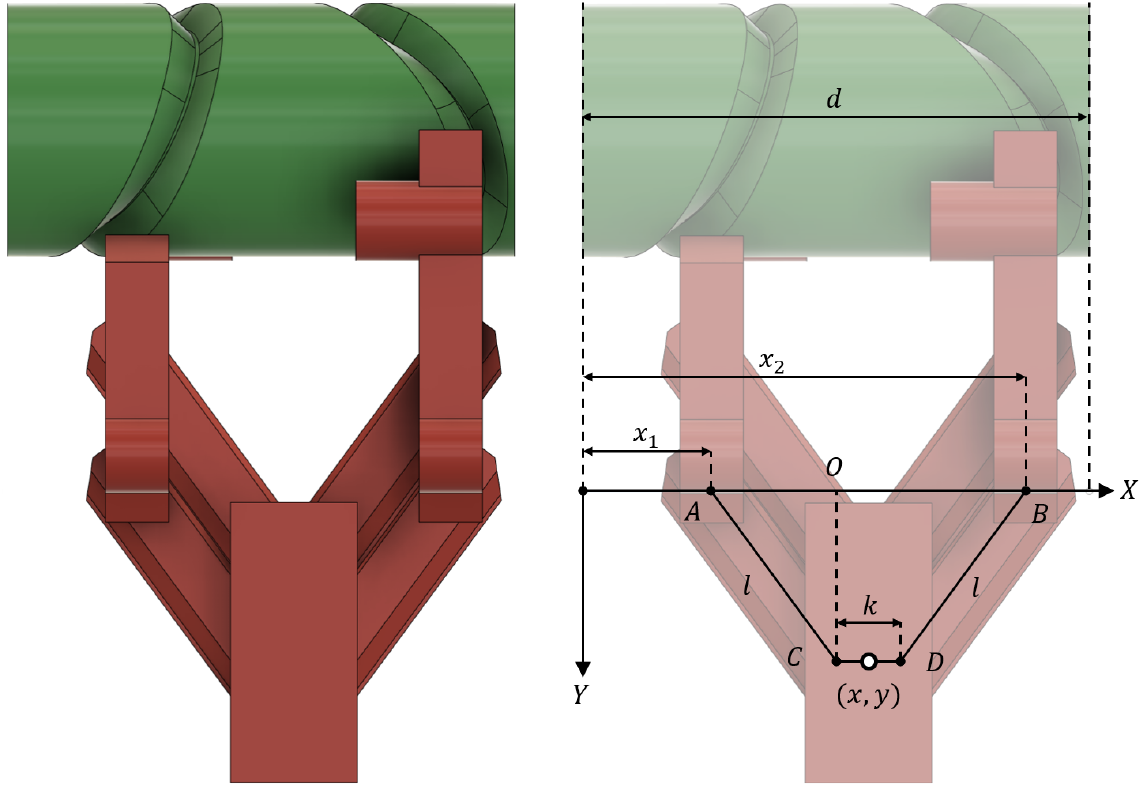
Diagram showing the inverse kinematics for the linkage, represented by the trapezoid ABCD.

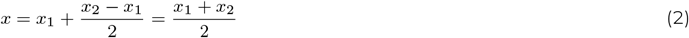

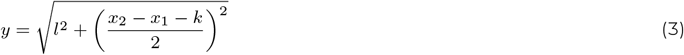

Rearranging the *x* equation, squaring the *y* equation and rearranging yields Equation (4) and Equation (5).

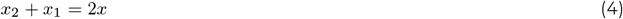

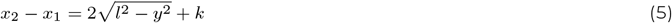

Summing the two equations and bisecting yields an expression for *x*_2_, Equation (6). Substituting its expression in *x*_2_ + *x*_1_ = *x* and rearranging leads to an expression of *x*_1_, Equation (7).

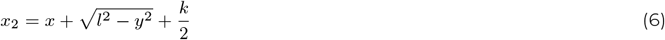

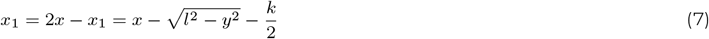

The position of the end effector was defined by the time *t*, and the desired coordinates *x*(*t*) and *y*(*t*) as functions of time. To get the grooves, we needed to substitute the coordinates in the inverse kinematics equations. An elliptical trajectory was chosen for the motion, as it is a trajectory with no discontinuities in its derivatives, and has been used for high-performance running robots ^91^. To obtain equations representative of the curves to be embossed on the surface of the cam, the ellipse was expressed in a parameterised form along the distance travelled across the cam surface, *s*. For *s* ∈ [0; 2*πr*_*c*_], a complete rotation is described. The ellipse equations parameterised around *s* are shown in Equation (8) and Equation (9). Here, *R*_*x*_, *R*_*y*_ represent the semi-major and semi-minor axes aligned with the *x*- and *y*-axes, respectively, and *x*_0_, *y*_0_ represent the offsets of the centre of the ellipse along the respective axes.

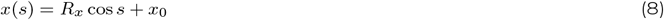

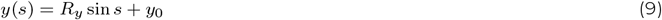

The properties of the ellipse most relevant to its use as a walking trajectory are the highest point along the *y*-axis *h*_*l*_, the lowest point *h*_*s*_, and the width *w*. The total width of the ellipse is just its major diameter, therefore 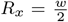. The minor axis would then be *h*_*s*_ − *h*_*l*_, therefore 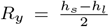. The vertical offset would be the average of the distances, 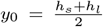. We desire that the ellipse centre be located on the midpoint of the cam horizontally, therefore 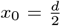, where *d* is the total cam length. Substituting these values for the parameters yields the following equations for the desired trajectory, Equation (10) and Equation (11):

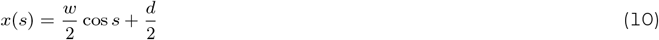

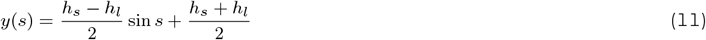

Substituting these equations in the inverse kinematics equations leads to the desired groove geometries, Equation (12) and Equation (13), across the cam surface, *s* ∈ [0; 2*πr*_*c*_].

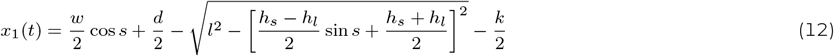

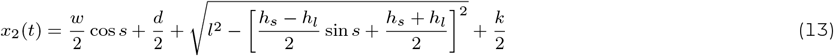

To obtain numerical expressions for these equations, the parameters *l, k, r*_*c*_, *w, h*_*s*_, *h*_*l*_ must be defined. *k* was assigned a value of 1.5 mm to match the thickness of the rings, allowing the mechanism to reach maximum extension when the rings touch. A smaller distance would lead to the mechanism failing to reach maximal extension at the maximum contraction of the mechanism, while a larger distance will require a longer cam to accommodate and is undesirable considering the space constraints imposed by the beetle body. *l* was set to 5mm due to manufacturing constraints, as this is the greatest length at which the linkage segments could be printed without warping when combined with a small brim. *r*_*c*_ was set to 3 mm, matching the radius of the motor. This leaves only the shape parameters of the ellipse to be defined, namely *w, h*_*s*_, *h*_*l*_. This design space has many solutions with varying feasibility for use in a walking machine. The desired properties of the mechanism include: (1) both the horizontal distance travelled and its speed were maximised whilst concurrently ensuring sufficient vertical distance to perform a step under loading - the vertical distance still nevertheless being made as small as possible as vertical motion does not result in forward motion, and it has to directly counter any payload on the robot (2) for a groove thickness *d*_*g*_, the groove was designed to be at least *d*_*g*_*/*2 distance from the ends of the cam and the other trajectories - this to prevent it from extending past the cam and coming into contact with the motor housing (3) all groove locations designed to be within the range of motion of the linkage, i.e. at no point should *y*(*s*) *> l* and (4) the slope of the cam was designed to be minimised to reduce wear and friction during operation. A numerical method was used to resolve the derivatives of the equations. Since the statements above can be expressed as inequalities, we used MATLAB’s `fmincon` constrained nonlinear optimisation function, which for a vector of variables *x*, minimises a given loss function *f* (*x*) while also obeying certain equalities *c*_*eq*_ (*x*) = 0, inequalities *c*(*x*) *<*= 0, and upper and lower bounds on the values of *x*. The loss function used with `fmincon` is −| (*w*)^*a*^ (*h*_*s*_ − *h*_*l*_)^*b*^|, representing a product of ellipse width and height, weighted by the parameters *a* and *b*. For the generation of the cam used in all experiments, we used *a* = 1.1 and *b* = 1.

### Experimental mapping of trajectories

Due to its small size and accuracy requirements, the proposed cam mechanism is at the limit of what can be manufactured with a conventional FDM 3D printer. This makes it particularly sensitive to manufacturing tolerances and defects. As such we conducted an experiment to confirm that the trajectory of the leg tip is sufficiently accurate to its desired path, even when accounting for manufacturing-related geometrical distortions. The experiment setup consisted of the mechanism, a vertical clamp workbench, a Chronos 1.4 Monochrome slow-motion camera with a Computar 12.5-75mm f/1.2 zoom lens on a tripod, a halogen lamp and an Energiser Vision HD headlight. The mechanism was clamped by the motor using the workbench-integrated table vice. The camera was placed 1m away from the demonstrator and the zoom lens was set to a focal distance of 75mm, to minimise perspective distortion. The camera shutter was left completely open to maximise the available light. The two lamps were chosen to prevent flickering due to AC current - the halogen lamp circumvents this as the heated tungsten filament does not change in light, and the LED torch is continuously powered by DC current and therefore does not flicker.

Eight cams and linkages were manufactured in a single print to prevent irregularities stemming from build plate levelling or printer wear. All parts were manufactured in the same orientation, and the location of the Z-seam was normalised by setting it to a consistent location, thus ensuring that G-code operations were the same for all parts. After manufacturing, each mechanism was actuated for at least 1 minute before testing to remove any transient behaviour stemming from printer irregularities such as surface imperfections caused by stringing. During the experimental phase, a cam-linkage pair was attached to a motor (Garosa 6mm Micro Motor) and recorded at a rate of 1057 frames per second. Each cam-linkage pair was tested for both the 4× and 2× gearbox, to ascertain whether the motive speed affects the trajectory. Each mechanism was recorded for a total of 8 s.

After recording, motion tracking was performed on the footage using the Tracker video analysis software ^92^. To compare between the experiments using the different motors, the 4× gearbox motor footage was sped up by a factor of the reduction ratio between the 4× and 2× gearboxes as reported by the manufacturer (26.43). For the 4 × gearbox experiments with 17469 frames, this results in 671 samples, and the same amount of frames were used for data collection from the 2× gearbox experiments. We used an auto-tracking operation on a dot applied with a permanent marker to the middle of the end effector. The scale of the footage was then calibrated to convert measurements from pixel units to millimetres. The motor housing was used for calibration, as it is a prominent feature in every video, and has a known height of 8.50 ± 0.05 mm, which was measured for each sample using vernier callipers. After tracking, the trajectory of points was imported into MATLAB for processing. Since the coordinate system of Tracker was set to begin at the top left corner of the video, the resulting trajectory of the point needed to be offset and centered to com-pare it against the theoretical elliptical trajectory. This was achieved by subtracting offsets *o*_*x*_, *o*_*y*_ from the coordinates of every point, assuming **p**_**x**_, **p**_**y**_ are vectors containing the respective Cartesian coordinates of all points in a trajectory. The offsets were then evaluated as the averages of the closest and farthest points in the Cartesian directions, Equation (14) and Equation (15):

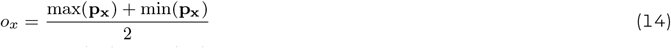

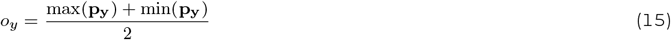

### Payload experimentation

A payload capacity experiment was conducted on Poka to compare its performance against that of *Xylorctes thestalus*. A silicone mat was laid on a flat surface, and two markers were placed 5 cm apart (measured with an accuracy of 0.05mm), defining the analysed distance over which the necro-robot, Poka, would travel. A 650 ml water bottle was used to exert a variable payload. The bottle was chosen as its weight could be varied by filling it with water, while the geometry of the payload would remain consistent, which decreased the chances of confounding variables arising through testing. A cylindrical water bottle holder was manufactured to secure the bottle to the necro-robot, Poka, and this was attached to the beetle using cyanoacrylate glue. The holder was positioned above the approximate centre of mass of Poka, which was estimated by edge-balancing the beetle. Poka’s legs were aligned with the edge of one of the insulation tape markers at the start of experimentation. The water bottle was then filled to achieve a target weight, and Poka was activated. The necro-robot was recorded using a camera (Monitech 4K Ultra HD, Monitech Ltd, Staffordshire, UK), and timed over its ambulation duration over the 5 cm distance. The experiment was conducted eight times (n = 8) at each weight tested. The payload weight was increased until the necro-robot was unable to locomote. We additionally recorded the current consumption for different payload weights used. The current draw can be used to estimate the power exerted per unit body and total weight, enabling a direct performance comparison with that of *Xyloryctes thestalus*. Furthermore, evaluating the current draw when the mechanism is not under load allows an estimation of friction within the system, which in turn enables the determination of the efficiency of the mechanism. A Keysight EDU34450A digital multimeter (Keysight Technologies Netherlands B.V., Amsterdam, Netherlands) was set to current measuring mode, and connected in series to the motor. The setup was powered at 3.3V, the rated voltage of the motors (using the Keysight EDU34450A powersupply). The multimeter was to a laptop and operated using the Keysight BenchVue Digital Multimeter interface for data logging. The multimeter was set to a sample rate of 25 Hz.

## Supporting information

Zip file containing all supplementary materials

## ELECTRONIC SUPPLEMENTARY MATERIALS

The following supplementary videos are available:

- Electronic Supplementary Video 1: Slow motion video of trajectory analysis using a 2× gear
- Electronic Supplementary Video 2: Poka carrying a payload of 500 g, which is an equivalent payload ratio of 6847%

The following supplementary sections are available within the Electronic Supplementary Material file:

- Proof of Cam Accuracy
- Statistical Analysis on the Interchangeability of the Cam-Linkage Couplings
- Individual Trajectories Mapped against the Ellipses for both 2× and 4× Motors
- Current Data Increasing Linearly with Payload
- Tabulated Payload Ratios of Poka Compared against Numerous Robots
- Separated Body Parts of the Five Horned Rhinocerous Beetle

## DATA AVAILABILITY

Data for this publication will be made available through Edinburgh DataShare (https://datashare.ed.ac.uk/) and can also be made available from the corresponding author on request.

## AUTHOR CONTRIBUTIONS

Conceptualization (PA); Data curation (YT); Formal analysis (YT, PA); Funding acquisition (na); Investigation (YT, PA); Methodology (YT, PA); Project administration (PA); Resources (YT, PA); Software (na); Supervision (PA); Validation (YT, PA); Visualisation (YT); Roles/Writing - original draft (YT); Writing - review and editing (PA).

## OPEN ACCESS STATEMENT

For the purpose of open access, the authors have applied a Creative Commons Attribution (CC BY) license to any author accepted manuscript version arising from this submission.

