## Supplementary material for "Poka: a necro-robot beetle with a measured payload ratio of 6847 %": Zip file containing all supplementary materials: Electronic Supplementary Material_compressed.pdf

### PROOF OF MECHANISM ACCURACY

The accuracy of the mechanism manufactured for Poka can be compared against other 3D-printed cam mechanisms. Cheng et al.[1], detail a computational method that can be used to generate a variety of cam-follower mechanisms. One of the mechanisms reported, transforms rotary motion into 2D Cartesian motion, fulfilling the same functional requirement as the proposed mechanism. As the paper does not provide numerical estimations for the accuracy of the trajectory, we extracted data points with Cartesian coordinates from the pictures using WebPlotDigitiser[2]. After performing the analysis, we note that the mechanism by Cheng et al. possesses a mean error of 1.8 mm across 72 sampled points. This error is ca. 25-fold larger than the mean error for the 4× gearbox motor (0.0789 mm) and ca. 19-fold larger than the 2× gearbox motor (0.0964). When considering the limited accuracy of FDM 3D printing for milimetre-scale designs, alongside the fact that the mechanism successfully enables the necrobot to walk, this accuracy is considered as being more than satisfactory. The comparative trajectories are shown in Figure 1.

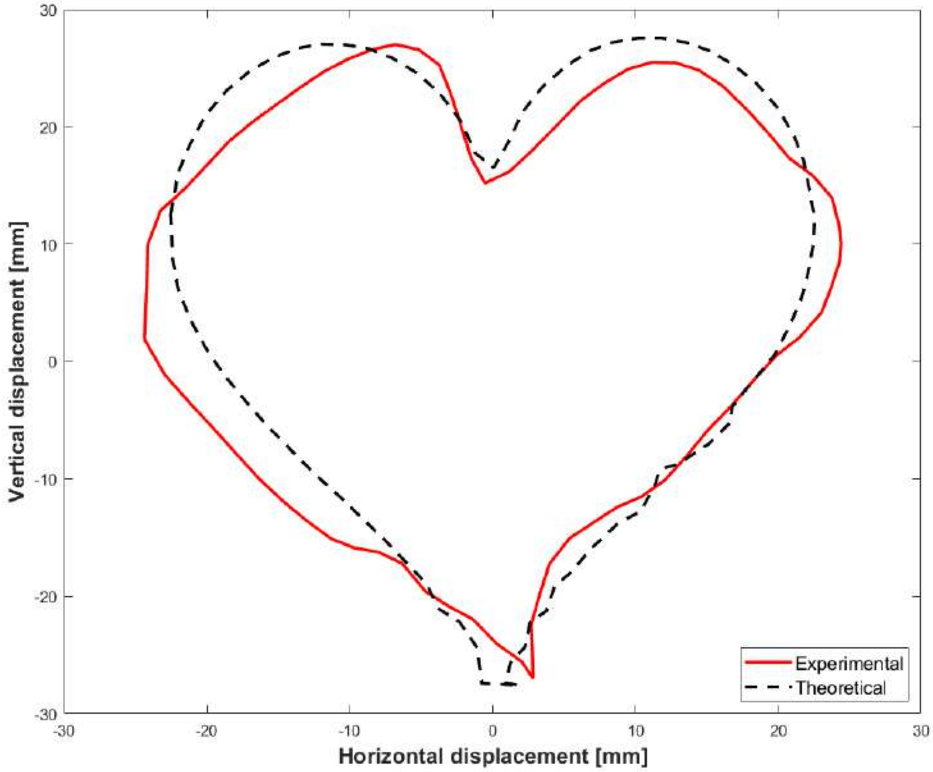


**Figure 1:** Visualisation of experimental trajectory (red) versus desired trajectory (black) of Cheng et al. mechanism[1].

### STATISTICAL ANALYSIS ON THE INTERCHANGEABILITY OF THE CAM-LINKAGE COUPLINGS

To ascertain the interchangeability (and thus relative repeatability) of the cam-linkage couplings, we collected data for two different sample sets. The first sample set comprised randomly paired cam-linkage couplings, while in the second sample set, we paired all cams with the same single linkage. The data collected enables statistical comparison, in order to determine the factors that might lead to statistically-significant differences in performance. We additionally wanted to check if the choice of motor affects the trajectory accuracy in these two sample sets, resulting in a total of three tests. To conduct the analysis, we used the Kruskal-Wallis test (`kruskalwallis` in MATLAB). The reason we used this test instead of more common statistical tests such as the analysis of variance (ANOVA) or Student's t-test is that it does not assume a normal distribution of the data [3]. A check for whether the data is normal or not, is possible through a one-sample Kolmogorov-Smirnov test [4] on each of the trajectories output, using the MATLAB function `kstest`. The p-value for all trajectories is smaller than $1 \times 10^{-10}$, suggesting that the normality hypothesis can be strongly rejected.

**Table 1:** Results of statistical significance analysis performed using the Kruskal-Wallis test.

| Sample set 1 | Sample set 2 | p-value | p $< 0.05$? |
|---|---|---|---|
| 2x gearbox, random cam and linkage | 2x gearbox, random cam only | 0.1033 | True |
| 4x gearbox, random cam and linkage | 4x gearbox, random cam only | 0.4945 | True |
| 2x gearbox, all data | 4x gearbox, all data | 0.0143 | False |

The results suggest that despite a small variance in the mean, the speed of the motor does not affect the mean accuracy in a statistically-significant way, while the randomly paired cam-linkage couplings leads to a statistically-significant results. As the mean errors are larger in the case of randomly paired cam-linkage couplings, Table 1, it can be suggested that the linkage is able to significantly affect the behaviour of the cam mechanism. This could be related to defects, which in turn may provide explanation for the trajectory errors, since the linkage is composed of small parts which will be dimensionally affected by the presence of defects.

#### INDIVIDUAL TRAJECTORIES MAPPED AGAINST THE ELLIPSES FOR BOTH 2$\times$ AND 4$\times$ MOTORS

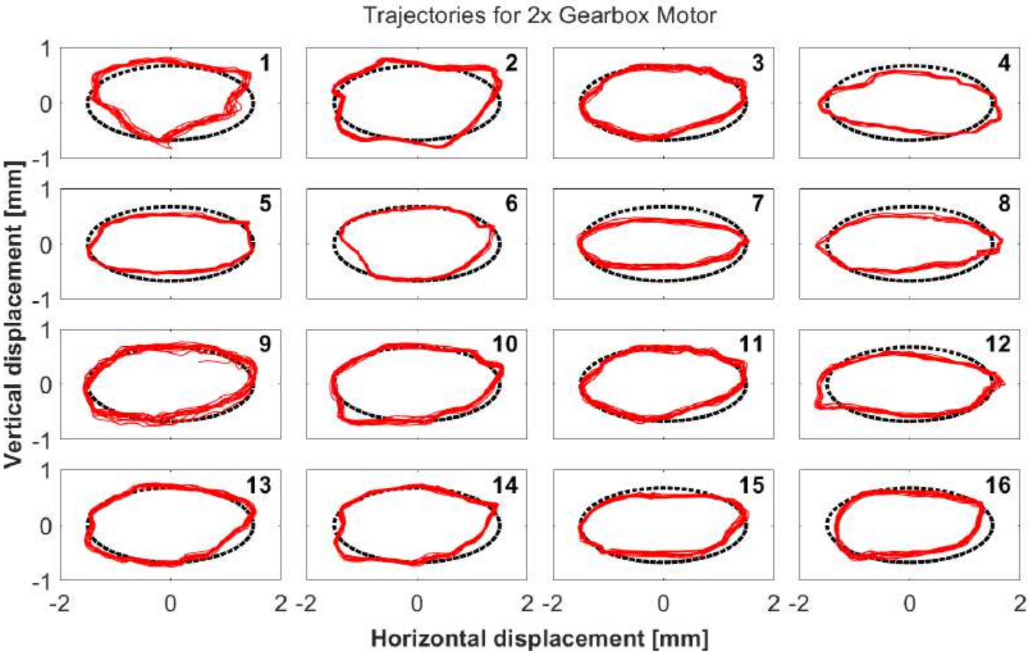


A 2x gearbox motor

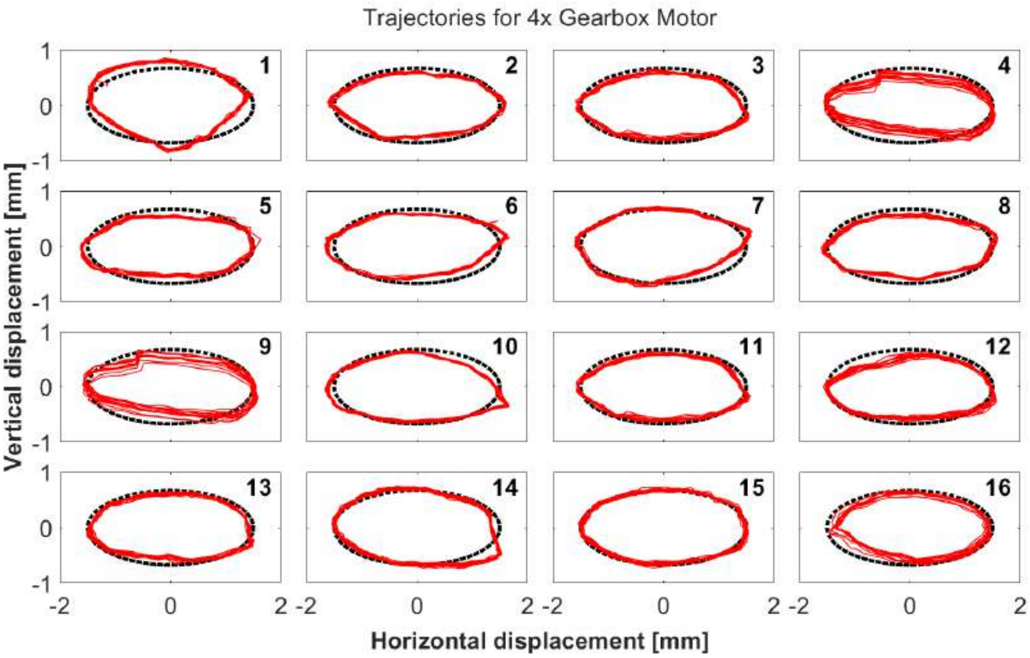


B 4x gearbox motor

**Figure 2:** Visualisation of all 32 experimental trajectories (red) overlaid on the desired theoretical response (black).

**CURRENT DATA INCREASING LINEARLY WITH PAYLOAD**

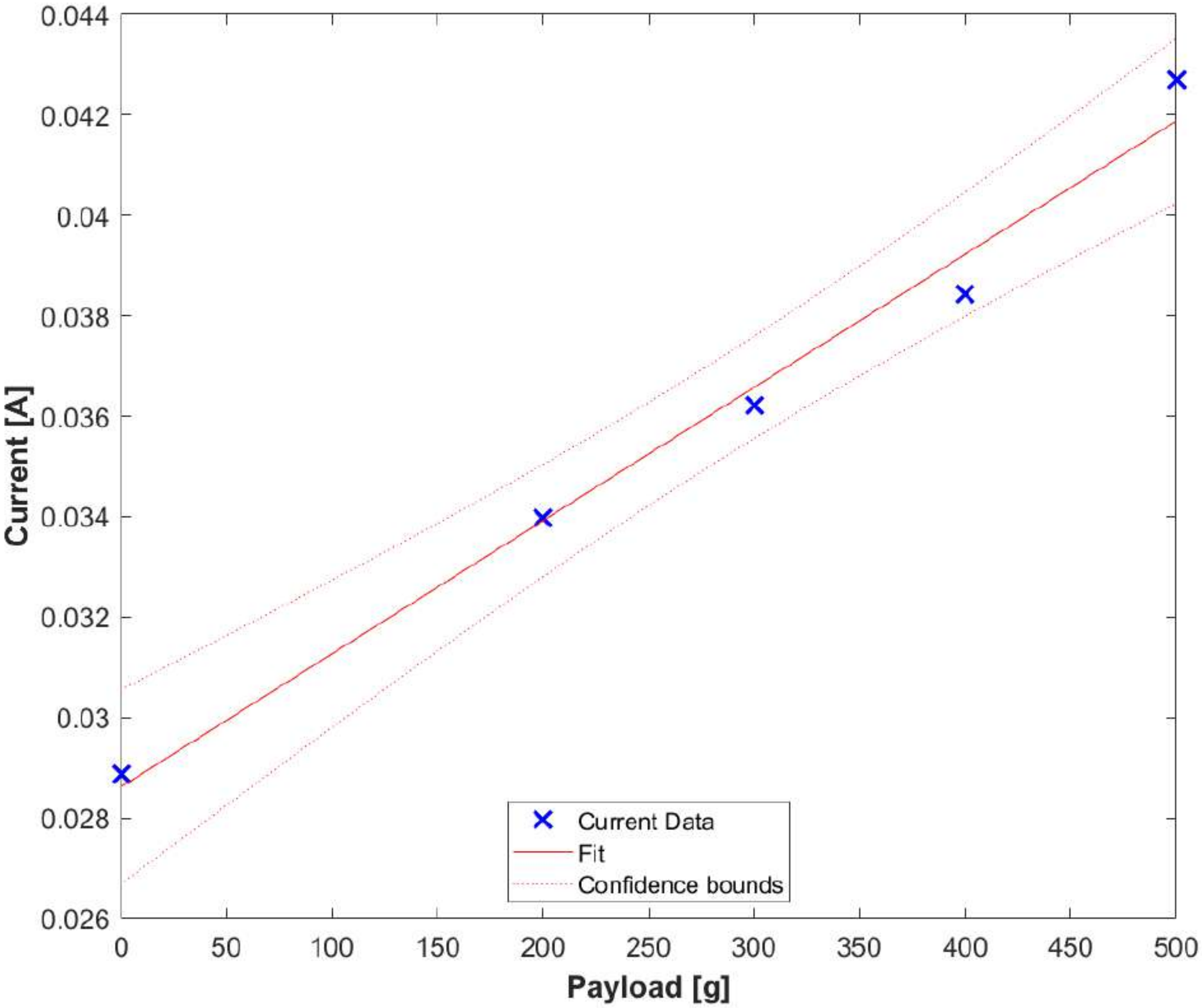


**Figure 3:** Mean current data with a linear regression model fit. The current data scales in a highly linear fashion as the load increases, with $R^2 = 0.9856$.

**TABULATED PAYLOAD RATIOS OF POKA COMPARED AGAINST NUMEROUS ROBOTS**

**Table 2:** Weight, maximum payload carried, and payload ratio all robots referred to in Figure 15 of the paper.

| Name | Weight [kg] | Payload Ratio [kg/kg] | Source |
|---|---|---|---|
| Poka | 0.0073 | 68.47 | |
| Baby Elephant | 130 | 0.7692 | [5] |
| Anymal D | 50 | 0.3 | [6] |
| Unitree A1 | 12 | 1 | [7] |
| BigDog | 109 | 1.41 | [8] |
| SpotMini | 30 | 0.47 | [9] |
| Kirin | 50 | 2.5 | [10] |
| SuperBot | 2.636 | 5.3 | [11] |
| RHex | 7 | 1.134 | [12] |
| Mini-Whegs | 0.146 | 2 | [13] |
| Minitaur | 5 | 0.4 | [14] |
| Kang et al. | 30 | 0.66 | [15] |
| StarlETH | 23 | 0.16 | [16] |
| PlusTech Walking Machine | 3400 | 0.556 | [17] |
| Tri-ATHLETE | 720 | 0.2577 | [18] |
| Landmaster (Electric) | 82 | 0.2679 | [19] |
| Landmaster (Hydraulic) | 3950 | 0.2020 | [19] |

**SEPARATED BODY PARTS OF THE FIVE HORNED RHINOCEROS BEETLE**

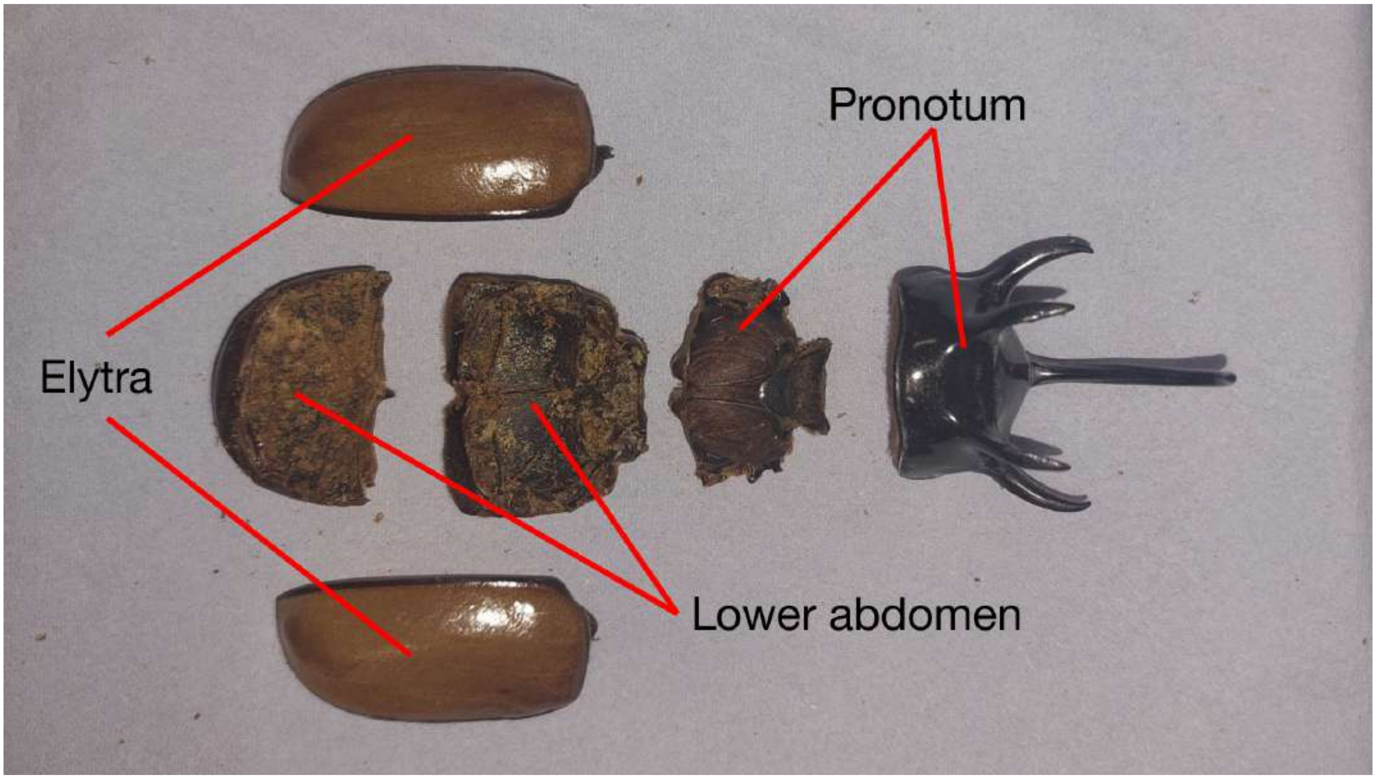


**Figure 4:** Body parts and positions of the five horned rhinoceros beetle showing specifically the elytra, lower abdomen, and protonum.

[10] Yueheng Zhou, Ming Liu, Chaoyang Song, and Jianwen Luo. Kirin: A quadruped robot with high payload carrying capability, 2022.

[18] Matt Heverly, Jaret Matthews, Matt Frost, and Chris Quin. Development of the tri-athlete lunar vehicle prototype. In *Proceedings of the 40th Aerospace Mechanisms Symposium*, page 20100021930, 2010.

[19] Hongchao Zhuang, Jiaju Wang, Ning Wang, Weihua Li, Nan Li, Bo Li, and Lei Dong. A review of footterrain interaction mechanics for heavy-duty legged robots. *Applied Sciences*, 14(15), 2024.
